## Supplementary Discussions, Tables, Legends and Descriptions of Supplementary Data for "Widespread potential for phototrophy and convergent reduction of lifecycle complexity in the dimorphic order *Caulobacterales*"

Version: 2025-04-25

SUPPLEMENTARY DISCUSSIONS 1

Supplementary Discussion 1 | Taxonomic descriptions according to the International Code of Nomenclature of Prokaryotes (ICNP) 1

Description of *Aquidulcibacteraceae* fam. nov. 1

Emended description of the genus *Aquidulcibacter* Cai *et al*. 2017 1

Emended description of the genus *Pseudaquidulcibacter* Liu *et al*. 2022 1

Emended description of the genus *Vitreimonas* Asem *et al*. 2020 2

Description of *Vitreimonas silvestris* comb. nov. 2

Description of *Poindextera* gen. nov. 2

Description of *Poindextera montana* comb. nov. 2

Description of *Oceanicaulis satelles* comb. nov. 3

*Hyphomonas neptunia* corrig. (Leifson 1964) Moore *et al*. 1984 (syn. *Hyphomonas hirschiana* Weiner *et al*. 1985) 3

Supplementary Discussion 2 | Additional taxonomic notes 4

Motivation for the description of *Aquidulcibacteraceae* fam. nov. 4

*Terricaulis silvestris* Vieira *et al*. 2020 belongs to the genus *Vitreimonas* Asem *et al*. 2020 4

*Phenylobacterium montanum* Tang *et al*. 2025 does not belong to the genus *Phenylobacterium* Lingens *et al*. 1985 5

*Alkalicaulis satelles* Kevbrin *et al*. 2021 belongs to the genus *Oceanicaulis* Strömpl *et al*. 2003 6

*Hyphomonas hirschiana* Weiner *et al*. 1985 belongs to the species *Hyphomonas neptunia* corrig. (Leifson 1964) Moore *et al*. 1984 6

Additional taxonomic notes 6

Supplementary Discussion 3 | Phylogeny, ecology, and physiology of *Acaudatibacter* gen. nov. 8

Phylogeny of *Acaudatibacter* gen. nov. 8

Ecology and putative physiology of *Acaudatibacter* gen. nov. 9

Ecology and physiology of *Ac. aquilonius* sp. nov., *Ac. boreus* sp. nov., and *Ac. lapponiensis* sp. nov. 10

Supplementary Discussion 4 | Taxonomic descriptions according to the SeqCode code of nomenclature 13
Description of *Acaudatibacter* gen. nov. 13
Description of *Acaudatibacter aquilonius* sp. nov. 14
Description of *Acaudatibacter boreus* sp. nov. 15
Description of *Acaudatibacter lapponiensis* sp. nov. 16

SUPPLEMENTARY FIGURES 18

SUPPLEMENTARY TABLES 24

SUPPLEMENTARY VIDEOS 27

SUPPLEMENTARY DATA 28

REFERENCES FOR SUPPLEMENTARY INFORMATION 32

### SUPPLEMENTARY DISCUSSIONS

#### Supplementary Discussion 1 | Taxonomic descriptions according to the International Code of Nomenclature of Prokaryotes (ICNP)

Based on our phylogenomic analyses, we make the following taxonomic proposals according to the International Code of Nomenclature of Prokaryotes (ICNP). Motivations for these proposals are found in **Supplementary Discussion 2**.

##### Description of *Aquidulcibacteraceae* fam. nov.

*Aquidulcibacteraceae* (*A.qui.dul.ci.bac ter.a.ce'ae*. N.L. masc. n. *Aquidulcibacter*, type genus of the family; -*aceae* ending to denote a family; N.L. fem. pl. n. *Aquildulcibacteraceae* the family of the genus *Aquidulcibacter*).

Gram-negative, rod-shaped bacteria. Do not form spores. Motile by means of a single polar flagellum. Some species form one or multiple prosthecae. Reproduce by binary fission or by prosthecal budding. Some species form rosettes. Colonies are circular, and white, yellow, or pink. Chemotrophic or phototrophic heterotrophs. Strict or facultative aerobes. Grow optimally in absence of NaCl. Q-10 is the major respiratory quinone. Members of the family can be isolated from cyanobacterial aggregates, soil, or water treatment facilities, and occur in algal consortia and freshwater. The G+C content range is 41.4–63.5%. Currently, the family comprises the type genus *Aquidulcibacter* Cai *et al.* 2017[1], and the genera *Pseudaquidulcibacter* Liu *et al*. 2022[2] and *Vitreimonas* Asem *et al*. 2020[3]. Belongs to the order *Caulobacterales* and the class *Alphaproteobacteria*.

##### Emended description of the genus *Aquidulcibacter* Cai *et al.* 2017

The description is as given by Cai *et al*. 2017[1] with the following amendment. Phylogenetically belongs to the family *Aquidulcibacteraceae*.

##### Emended description of the genus *Pseudaquidulcibacter* Liu *et al.* 2022

The description is as given by Liu *et al*. 2022[2] with the following amendment. Phylogenetically belongs to the family *Aquidulcibacteraceae*.

**Emended description of the genus *Vitreimonas* Asem *et al.* 2020**

The description is as given by Asem *et al.* 2020[3] with the following amendment. Cells are motile by means of a single flagellum and produce one or multiple prosthecae. Some members form branched prosthecae. Cells reproduce by binary fission or by prosthecal budding. Catalase and oxidase reactions, as well as major fatty acid profiles, vary between species. Polar lipids include glycolipids. Phylogenetically belongs to the family *Aquidulcibacteraceae*.

**Description of *Vitreimonas silvestris* comb. nov.**

Basonym: *Terricaulis silvestris* Vieira *et al.* 2020[4]. *Vitreimonas silvestris* (*sil.ves'tris.* L. fem. adj. *silvestris*, referring to the forest soil from which the type strain was isolated) is closely related to *Vitreimonas flagellata* according to phylogenomics and genomic sequence similarity, but is different from this species by morphology and mode of cell division[3, 4]. Differs from *Vitreimonas flagellata* in the oxidase and catalase tests, by its fatty acid profile, and by its salt tolerance. Further distinguished from *Vitreimonas flagellata* by its lack of activities for the enzymes α-chymotrypsin, esterase (C4), esterase lipase (C8), α-fucosidase, naphthol-AS-BI phosphohydrolase, and trypsin. The type strain is 0127_4$^{T}$ (= DSM 104635$^{T}$ = CECT 9243$^{T}$).

**Description of *Poindextera* gen. nov.**

*Poindextera* (*Poin.dex'te.ra*. N.L. fem. n. *Poindextera*, named after Jeanne S. Poindexter, who contributed greatly to the study of the *Caulobacteraceae*).

Gram-negative, rod-shaped bacteria. Do not form spores. Reproduce by binary fission. Colonies are circular, convex, and unpigmented. Aerobic, chemotrophic, mesophilic. Catalase-negative and oxidase-positive. The major respiratory quinone is Q-10. The major fatty acids are $C_{18:1}$ ω7c and $C_{16:0}$. Phosphatidylglycerol, three unidentified glycolipids, and one unidentified phosphoglycolipid are the major polar lipids. Belongs to the family *Caulobacteraceae* in the class *Alphaproteobacteria*. The type species is *Poindextera montana*.

**Description of *Poindextera montana* comb. nov.**

Basonym: *Phenylobacterium montanum* Tang *et al.* 2024[5]. *Poindextera montana* (*mon.ta'na.* L. fem. adj. *montana*, of a mountain) is phylogenetically distinct from members of the genera *Phenylobacterium* and *Caulobacter*. The description is as given by Tang *et al.* 2024[5]. The type strain is S6$^{T}$ (= NBRC 115419$^{T}$ = GCMCC 1.18594$^{T}$).

**Description of *Oceanicaulis satelles* comb. nov.**

Basonym: *Alkalicaulis satelles* Kevbrin *et al.* 2021[6]. *Alkalicaulis satelles* is closely related to *Oceanicaulis alexandrii* according to genomic sequence similarity, but is different from this species in the oxidase test, by its fatty acid profile, and by its genetic potential for type II anoxygenic phototrophy. Further distinguished from *Oceanicaulis alexandrii* by its lack of activities for the enzymes lipase (C14), acid phosphatase, and cystine arylamidase, and inability to utilize lactose, lactulose, and maltose. The type strain is G-192$^T$ (= KCTC 72746$^T$ = VKM B-3306$^T$).

***Hyphomonas neptunia* corrig. (Leifson 1964) Moore *et al.* 1984 (syn. *Hyphomonas hirschiana* Weiner *et al.* 1985)**

The description is based on the descriptions of *Hyphomonas neptunia* corrig. (Leifson 1964)[7] Moore *et al.* 1984[8] and *Hyphomonas hirschiana* Weiner *et al.* 1985[9]. The description of the species *Hyphomonas neptunia* corrig. was given by Moore *et al.* (1984)[8]. Strain VP5$^T$ (= ATCC 33886$^T$ = CIP 106773$^T$ = DSM 5152$^T$), the type strain of *Hyphomonas hirschiana*, is a reference strain of *Hyphomonas neptunia*. The type strain is strain 14-15$^T$ (= ATCC 15444$^T$ = BCRC 10690$^T$ = CCRC 10690$^T$ = DSM 5154$^T$ = IFAM LE-670$^T$ = IFO 14232$^T$ = LE 670$^T$ = NBRC 14232$^T$ = NCIMB 2023$^T$).

#### Supplementary Discussion 2 | Additional taxonomic notes

Based on our phylogenomic analyses, we make the following taxonomic notes. Formal taxonomic proposals are found in **Supplementary Discussion 1**.

**Motivation for the description of *Aquidulcibacteraceae* fam. nov.**

In phylogenies of concatenated single-copy marker genes conserved among *Alphaproteobacteria*, type strains of the recently described monotypic genera *Aquidulcibacter*[1], *Pseudaquidulcibacter*[2], and *Terricaulis*[4] assigned to the family *Caulobacteraceae*, as well as *Vitreimonas*[3] assigned to the family *Hyphomonadaceae*, consistently form a cluster separate from other members of *Caulobacteraceae* and *Hyphomonadaceae* (**Fig. 1a**, **Supplementary Figs. S1** and **S2**). This cluster also includes "*Candidatus* Phycosocius bacilliformis"[10, 11] and "*Candidatus* Viadribacter manganicus"[12] and is placed with good support (98% non-parametric bootstrap proportions [npBP] sister to *Hyphomonadaceae* (**Supplementary Fig. S1**). These results are in line with the current Genome Taxonomy Database (GTDB) taxonomy[13] (R220), which assigns these taxa into the "f__TH1-2" placeholder family separate from *Caulobacteraceae* and *Hyphomonadaceae* (named after *Aquidulcibacter paucihalophilus* TH1-2$^{T}$, the earliest validly described member of the group). We therefore propose the classification of this cluster as family *Aquidulcibacteraceae* fam. nov., as described in **Supplementary Discussion 1**. Notably, the placement of *Aquidulcibacteraceae* fam. nov. sister to *Hyphomonadaceae* is perhaps also congruent with the prosthecate budding mode of reproduction of the *Aquidulcibacteraceae* species *Vitreimonas silvestris* comb. nov. (*Terricaulis silvestris*[4]), a trait characteristic of *Hyphomonadaceae* that is not found in *Caulobacteraceae* sensu stricto (**Fig. 1a**).

***Terricaulis silvestris* Vieira *et al.* 2020 belongs to the genus *Vitreimonas* Asem *et al.* 2020**

*Terricaulis silvestris* 0127_4$^{T}$ [4] and *Vitreimonas flagellata* SYSU XM001$^{T}$ [3] are closely related and should be considered separate species of the same genus (**Supplementary Figs. S1–S2**), given their ANI of 79.6% below the 95% species cutoff[14, 15] (**Supplementary Data S11**) and their high AAI of 74.7% above the 65% genus cutoff[14, 16] (**Supplementary Data S12**). Consistent with this, in GTDB taxonomy releases R202 and R207, they were both classified as genus *Terricaulis*, and in later releases R214 and onward they have been reclassified as genus *Vitreimonas*. Notably, genomic AAI scores also place "*Candidatus* Viadribacter manganicus"[12] within this genus (**Supplementary Data S12**). Given the earlier description of genus *Vitreimonas*[3] than *Terricaulis*[4], we propose the reclassification of *Terricaulis silvestris* as *Vitreimonas silvestris* comb. nov. (type strain 0127_4$^{T}$ = DSM 104635$^{T}$ = CECT 9243$^{T}$), and the amendment of *Vitreimonas* Asem *et al.* 2020[3], as described in **Supplementary Discussion 1**.

#### *Phenylobacterium montanum* Tang *et al.* 2025 does not belong to the genus *Phenylobacterium* Lingens *et al.* 1985

In our phylogenomic analyses, the recently described species *Phenylobacterium montanum*[5] consistently clusters in the family *Caulobacteraceae*, but outside of its assigned genus *Phenylobacterium* and its sister genus *Caulobacter* (**Supplementary Figs. S1** and **S2**, **Extended Data Fig. 4**). Consistent with this phylogenetic placement, pairwise AAI comparisons of *Phenylobacterium* and *Caulobacter* genomes to *P. montanum* S6$^{T}$ are very similar between the two genera and are notably around the proposed 65% genus cutoff[14, 16]; for *Phenylobacterium* (n = 41) AAIs range between 64.77–66.63% and *Caulobacter* (n = 43) they range between 64.74–66.26% (**Supplementary Data S12**). Additionally, in the GTDB release R207 and onwards, the genome of strain S6$^{T}$ has been assigned to the uncharacterized genus-level clade “BOG-935”, instead of *Phenylobacterium* or *Caulobacter*. Thus, given that strain S6$^{T}$ is phylogenetically distinct from the genera *Phenylobacterium* and *Caulobacter*, which otherwise comprise its closest described relatives, this strain represents a new genus within the family *Caulobacteraceae*. For this, we propose the name *Poindextera* gen. nov., and the transfer of *Phenylobacterium montanum* Tang *et al.* 2025[5] to this genus as *Poindextera montana* comb. nov. (descriptions in **Supplementary Discussion 1**).

Additionally, we note that strain S6$^{T}$ was reported as non-motile under the studied growth conditions[5]. However, strain S6$^{T}$ has genetic potential for flagellar motility, including extensive suites of flagellar, chemotaxis, and dimorphic development genes (**Fig. 2b**, **Supplementary Fig. S4b–d**). Additionally, it has genetic potential for type IV pili and polar holdfast adhesin (**Fig. 2b**). Taken together, the genetic potential of strain S6$^{T}$ suggests that it has a dimorphic developmental program. Moreover, consistent with its gray, pale colonies on modified SSE/HD agar[5], strain S6$^{T}$ lacks carotenoid biosynthesis genes (**Supplementary Fig. S9b,c**).

##### ***Alkalicaulis satelles* Kevbrin *et al.* 2021 belongs to the genus *Oceanicaulis* Strömpl *et al.* 2003**

The genome of *Alkalicaulis satelles* G-192$^{T}$ [6] has an AAI to *Oceanicaulis alexandrii* DSM 11625$^{T}$ [17] of 69.7% (**Supplementary Data S12**), placing it in the genus *Oceanicaulis* (**Supplementary Figs. S1–S2**). This conclusion is shared with current GTDB taxonomy (R220), and is further supported by its high (68.3%) percentage of conserved proteins compared to *O. alexandrii*[6] (above the proposed 50% genus cutoff[18]). We therefore propose the reclassification of *Alkalicaulis satelles* as *Oceanicaulis satelles* comb. nov. (type strain G-192$^{T}$ = KCTC 72746$^{T}$ = VKM B-3306$^{T}$), as described in **Supplementary Discussion 1**. Notably, *Oceanicaulis satelles* comb. nov. has genes for type II anoxygenic phototrophy (**Fig. 5b**, **Supplementary Fig. S10**)[19], genetic potential which it shares with multiple uncharacterized *Oceanicaulis* species (**Fig. 5b**, **Supplementary Fig. S10**).

##### ***Hyphomonas hirschiana* Weiner *et al.* 1985 belongs to the species *Hyphomonas neptunia* corrig. (Leifson 1964) Moore *et al.* 1984**

*Hyphomonas hirschiana* VP5$^{T}$ [9] and *Hyphomonas neptunia* 14-15$^{T}$ [7, 8] should be considered the same species given their ANI of ~100% (**Supplementary Data S1a**). This is likely not the result of genome mislabeling, as EMBOSS Needle alignment[20] of their Sanger-sequenced 16S rRNA genes (GenBank accessions KF863147.1 and KF863145.1, respectively) also reveals 100% 16S rRNA identity (not shown). Given priority of publication, *Hyphomonas hirschiana* is therefore to be considered a later heterotypic synonym of *Hyphomonas neptunia*. Thus, we propose the transfer of *H. hirschiana* strains to *H. neptunia*. An emended description of *Hyphomonas neptunia* is provided in **Supplementary Discussion 1**.

##### **Additional taxonomic notes**

We note that the genera *Litorimonas* and *Algimonas* of the *Maricaulaceae* family are paraphyletic (**Supplementary Figs. S1** and **S2**), with *L. cladophorae* clustering together with *Al. arctica*, sister to *L. taeanensis*. Given the earlier description of genus *Litorimonas*[21] than *Algimonas*[22], reclassification of *Algimonas arctica* as *Litorimonas arctica* comb. nov. (type strain KCTC 32513$^{T}$ = MCCC 1K00233$^{T}$ = SM1216$^{T}$) might be warranted—a conclusion consistent with current GTDB taxonomy. However, the genome of *Al. arctica* KCTC 32513$^{T}$ has only 65.2% and 65.5% AAI to *L. taeanensis* DSM 22008$^{T}$ and *L. cladophorae* KCTC

23968[T], respectively. We therefore do not formally propose taxonomic emendation of *Al. arctica*.

We further note that in comparison with *Aquidulcibacter paucihalophilus* TH1-2[T], "*Candidatus* Phycosocius bacilliformis" BOTRYCO-2[10, 11] has an AAI of 81.8% (**Supplementary Data S12**) and "*Candidatus* Phycosocius spiralis" BOTRYCO-1 has an AAI of 76.3%[23], placing them in the genus *Aquidulcibacter*. This conclusion is consistent with current GTDB taxonomy (R220). Thus, all members of the genus *Aquidulcibacter* with sequenced genomes have the genetic potential for type II anoxygenic photoheterotrophy (**Supplementary Fig. S9**)[23].

#### Supplementary Discussion 3 | Phylogeny, ecology, and physiology of *Acaudatibacter* gen. nov.

Here follow discussions on the biology and genomic characteristics of members of *Acaudatibacter* gen. nov. (GTDB taxon "g__Palsa-881") to provide context for their taxonomic description. Formal taxonomic proposals according to the SeqCode code of nomenclature are found in **Supplementary Discussion 4**.

##### Phylogeny of *Acaudatibacter* gen. nov.

Concatenated phylogenies of single-copy marker genes conserved among *Alphaproteobacteria* support the inclusion of *Acaudatibacter* gen. nov. within the family *Caulobacteraceae* (**Fig. 1a**, **Extended Data Fig. 4**, **Supplementary Figs. S1** and **S2**). The phylogenies place the genus *Acaudatibacter* gen. nov. as a close relative of the genera *Caulobacter* and *Phenylobacterium*. In phylogenies of the alphaproteobacterial genes included in GToTree[24], *Acaudatibacter* is placed as a sister group of the *Caulobacter–Phenylobacterium* clade (**Extended Data Fig. 4** and **Supplementary Fig. S2**). However, in a more robust tree inferred using 72 previously manually curated alphaproteobacterial genes[25], *Acaudatibacter* is instead placed sister to *Phenylobacterium* with good support (100% non-parametric bootstrap support) (**Supplementary Fig. S1**). Members of the genus have average amino acid identities (AAIs) to species genome representatives from other *Caulobacteraceae* genera that range between 58.0%–68.0% (**Supplementary Data S12**). Consistent with previously proposed genus delineation cutoff of 65% AAI[14, 16], AAIs between genomes of species belonging to the genus (as assigned by GTDB-Tk v2.1.1; GTDB taxonomy release R207) range between 65.4%–78.3%. AAIs between type genomes of species described here (*Ac. aquilonius*, *Ac. boreus*, and *Ac. lapponiensis*), range between 76.3%–78.3%. Thus, among *Acaudatibacter* gen. nov. species represented in our dataset, these three species are most closely related to each other, given that their genomes only have AAIs between 66.1%–72.5% to other species currently represented in the genus, and between 58.8%–66.1% to species of other *Caulobacteraceae* genera. This conclusion is further supported by phylogenomic analysis (**Extended Data Fig. 4**), their genomic repertoire, and their inferred similar physiology (see below). As a final note on their phylogeny, single-gene phylogenies inferred for PufM (**Supplementary Figs. S13–S14**) and BchY (**Supplementary Figs. S15–S16**) orthologs from *Ac. aquilonius*, *Ac. boreus*, and *Ac. lapponiensis* consistently place them as a distinct group that clusters with sequences from members of the genus-level clade "CAIMFV01", another clade of the *Caulobacter–Phenylobacterium* branch of the *Caulobacteraceae* family.

**Ecology and putative physiology of *Acaudatibacter* gen. nov.**

Sequences of the aquatic species *Ac. aquilonius*, *Ac. boreus*, *Ac. lapponiensis*, and *Ac.* sp. 23796 have been detected in metagenomes from boreal stratified freshwater bodies of Fennoscandia and Canada (**Fig. 6b**, **Supplementary Fig. S17**). In these habitats, *Ac. aquilonius*, *Ac. boreus*, *Ac.* sp. 23796 have been observed to reach particularly high relative abundances (~0.5%–1.5%) (**Fig. 6c**, **Extended Data Fig. 8b**). *Ac.* sp. SZAS AMP-5, another aquatic species recovered from wastewater (Shenzhen, China) (**Supplementary Fig. 6b**), also mapped to freshwater from the River Narmada in India (**Supplementary Fig. S17b**). Other species of the genus were detected in global terrestrial environments, including permafrost and glacier soil samples from the Arctic (Alaska, USA; Stordalen Mire and Tarfala, Sweden) and Antarctic (Mackay Glacier, Antarctica), and tropical rhizosphere soil (Minas Gerais, Brazil) (**Supplementary Fig. S5b**).

The genus includes two deep-branching species with genetic potential for flagellar motility, type IV pili (T4P), and holdfast adhesin production, and similar cell development regulation gene suites as *Caulobacter crescentus* (*vibrioides*) CB15 and other dimorphic *Caulobacteraceae* (**Fig. 2b**, **Extended Data Fig. 3**). Since the presence or absence of prostheca currently cannot be predicted from genome annotations alone, it is unclear whether *Acaudatibacter* gen. nov. includes prosthecate species. However, the largest clade of *Acaudatibacter* gen. nov. species (10/12 species) lack most genes for flagella, chemotaxis, and holdfast production, in contrast to the two deep-branching species. Since these putatively holdfast-lacking, non-flagellated species additionally lack a large number of cell development regulatory genes (**Extended Data Fig. 3**), mirroring the symmetrically reproducing apparently monomorphic species *Phenylobacterium immobile* (**Fig. 3**), these species may have monomorphic lifecycles. These putatively monomorphic *Acaudatibacter* species form a monophyletic group (**Extended Data Fig. 4**). Notably, unlike *P. immobile*, most putatively monomorphic *Acaudatibacter* gen. nov. species do encode T4P (**Extended Data Fig. 3**).

Like other *Caulobacteraceae* members, *Acaudatibacter* gen. nov. genomes contain signature genes for the Entner-Doudoroff Pathway, for respiration using an NADH-quinone oxidoreductase and ubiquinol-cytochrome *c* reductase (cytochrome $bc_1$), for an F-type ATPase, and for biosynthesis of polyphosphate and polyhydroxybutyrate granules (**Supplementary Fig. S9i** and **S11g**, **Supplementary Data S13**). As is also typical of *Caulobacteraceae*, most *Acaudatibacter* gen. nov. species have genes for high-affinity phosphate transporters (*pstABCS*) and phosphonate transporters (*phnDEC*), except rhizosphere species *Ac.* sp. 19682 and *Ac.* sp. 19683 which lack detectable *phnDEC* (**Supplementary Data S8** and **S13**).

#### Ecology and physiology of *Ac. aquilonius* sp. nov., *Ac. boreus* sp. nov., and *Ac. lapponiensis* sp. nov.

Metagenome assembled genomes (MAGs) of freshwater species *Ac. aquilonius*, *Ac. boreus*, and *Ac. lapponiensis* have been assembled from slightly acidic (pH ~5.0) freshwater samples, at water temperatures between 9.8–13.8°C, 9.5–16.1°C, and 7.4°C respectively (**Supplementary Fig. S5b**). These three species have been detected at high relative abundance in the upper anoxic layers of stratified freshwater bodies during the summer months, with *Ac. aquilonius* also reaching high relative abundance in oxic layers in some sample series (**Fig. 6c**). Such localization patterns indicate facultative anaerobic lifestyles, which is further supported by their genetic potential for aerobic respiration (**Supplementary Fig. S11g**). As further evidence of their potential adaptation to anoxic low-pH freshwater, genomes of *Ac. aquilonius*, *Ac. boreus*, and *Ac. lapponiensis* encode the ferrous iron transporter FeoB (**Supplementary Data S13**); in low-oxygen, low-pH environments, ferrous ($Fe^{2+}$) iron is abundant, while ferric ($Fe^{3+}$) iron is predominant under high-oxygen conditions[26]. At these upper anoxic strata, they likely photosynthesize using their genetic potential for type II anoxygenic photosynthesis (**Supplementary Fig. S11a–f**), as it typical of anoxygenic phototrophs[27]. This genetic potential includes a photosynthetic reaction center (RC), a light-harvesting I complex (LH1), and light-harvesting II complex (LH2), bacteriochlorophyll and carotenoid pigments, and a partial (*Ac. boreus*; potentially due to genome incompleteness, see **Results**) or complete (*Ac. aquilonius* and *Ac. lapponiensis*) Calvin-Benson-Bassham (CBB) cycle for carbon fixation (**Supplementary Fig. S12b**), which includes the accessory genes for red-type RuBisCO activase (*cbbX*) and XuBP phosphatase (*cbbY*) (**Fig. 5c**). Notably, like other *Caulobacterales* phototrophs (**Supplementary Fig. S9**), *Ac. aquilonius*, *Ac. boreus*, and *Ac. lapponiensis* lack the PufC photosynthetic reaction center cytochrome *c* subunit and the PufX reaction center protein present in some other *Alphaproteobacteria* (**Supplementary Figs. S10** and **S11e**).

Additionally, the non-homogenous distribution of *Ac. aquilonius*, *Ac. boreus*, and *Ac. lapponiensis* in the water column, suggests that they are able to regulate their buoyancy, despite lacking flagella. Whereas we could identify gas vesicle proteins (Pfam domains: 3×PF00741, 1×PF05120, 1×PF05121, 1×PF05800, 3×PF06386) in the phototroph *Rhodobacter sphaeroides* 2.4.1$^{T}$ (order *Rhodobacterales*), *Caulobacterales* genomes lacked such genes, indicating that these *Acaudatibacter* species possess other buoyancy regulation mechanisms, which could include T4P-mediated cell aggregation.

Based on KEGG Decoder, *Ac. aquilonius*, *Ac. boreus*, and *Ac. lapponiensis* have complete biosynthesis pathways for all standard amino acids, except valine and isoleucine. However, KEGG Decoder predicts these two pathways as incomplete for all *Caulobacterales* genomes, including the known amino acid prototrophs *Asticcacaulis excentricus* AC48$^{T}$ (CB 48$^{T}$)[28], *C. crescentus* CB15 [28], and *C. segnis* TK0059$^{T}$ [29] (**Supplementary Data S13**). The incorrect prediction stems from KEGG Decoder v1.3 requiring the presence of acetolactate synthase II small subunit IlvM [EC:2.2.1.6] (K11258) for these two pathways to be deemed complete (together with K00826, K01687, K00053, K01652, and K01653). However, this KEGG ortholog (KO) is absent from the vast majority of *Caulobacterales* genomes (**Supplementary Data S8** and **S9**). Instead, both large and small subunits of the corresponding enzyme [EC:2.2.1.6] are represented in most *Caulobacterales* genomes as K01652 and K01653. Therefore, since *Ac. aquilonius*, *Ac. boreus*, and *Ac. lapponiensis*—like the aforementioned known amino acids prototrophs—merely lack K11258, but otherwise have predicted complete amino acid pathways, we conclude that these *Acaudatibacter* gen. nov. species are likely amino acid prototrophs.

The genetic potential of *Ac. aquilonius*, *Ac. boreus*, and *Ac. lapponiensis* genomes differs on some points, as follows. *Ac. boreus* and *Ac. lapponiensis* have genetic potential for starch/glycogen biosynthesis and degradation, which *A. aquilonius* lacks (**Supplementary Data S13**). *Ac. boreus* and *Ac. lapponiensis* have genes for cytochrome *bd* terminal oxidase, which *A. aquilonius* lacks (**Supplementary Fig. S11g**). *Ac. aquilonius* and *Ac. boreus* encode thiosulfate dehydrogenase TsdA for thiosulfate oxidation, which *Ac. lapponiensis* lacks

(**Supplementary Data S13**). *Ac. boreus* has genetic potential for sulfide oxidation using the Sqr sulfide:quinone oxidoreductase (K17218), which *Ac. lapponiensis* and *Ac. aquilonius* lack (except for the low-quality *Ac. aquilonius* MAG “AM-lipid-02-D3_megahit_metabat_bin-0449”). Lastly, *Ac. boreus* and *Ac. lapponiensis* encode the SsuABC (K15553–K15555) sulfonate transporter, which *Ac. aquilonius* lacks.

#### Supplementary Discussion 4 | Taxonomic descriptions according to the SeqCode code of nomenclature

The SeqCode register list for *Acaudatibacter* gen. nov., *Acaudatibacter aquilonius* sp. nov., *Acaudatibacter boreus* sp. nov., and *Acaudatibacter lapponiensis* sp. nov. is available under the accession ID 9aocwnme (https://seqco.de/r:9aocwnme).

##### Description of *Acaudatibacter* gen. nov.

*Acaudatibacter* (A.cau.da.ti.bac'ter. Gr. pref. *a*-, not, without [inseparable prefix]; L. masc. adj. *caudatus*, tailed or having a tail; N.L. masc. n. *bacter*, rod; N.L. masc. n. *Acaudatibacter*, tailless rod).

Members of this genus have been identified in freshwater, wastewater, and in soils from permafrost active layers, of glacier regions, and of rhizospheres of *Barbacenia macrantha* and *Vellozia epidendroides*. ANI values among genomes representing separate species within the genus range between < 76.6% and 81.0%. AAI values among genomes representing separate species within the genus range between 65.4% and 78.8%. Genomes of this genus notably contain genes for the Entner-Doudoroff Pathway, for aerobic respiration using an NADH-quinone oxidoreductase, a ubiquinol-cytochrome *c* reductase, and a cytochrome *c* oxidase, an F-type ATPase, and for biosynthesis of polyphosphate and polyhydroxybutyrate. Some members have the genetic potential to produce bacteriochlorophyll and/or carotenoid pigments. Most members encode genes for type IV tight-adhesion pili, with some members further encoding genes for flagellar motility, chemotaxis, and holdfast formation, with genetic potential similar to members of the *Caulobacteraceae* family with obligate dimorphic lifecycles. However, most members likely have a monomorphic cell developmental program, as inferred from their lack genes for flagella, chemotaxis, and polar holdfast adhesin, as well as absence of a large number (> 20) of cell cycle regulation and polar morphogenesis genes present in related dimorphic taxa. The genus includes genomes with chemoheterotrophic, photoheterotrophic, and photoautotrophic genetic potentials. The taxon is supported as a genus-level group by phylogenomics and AAI, and corresponds to the GTDB taxonomy (R220) genus "g__Palsa-881".

The nomenclatural type for the genus is *Acaudatibacter aquilonius* GCA_903872075.1$^{Ts}$.

SeqCode URL: https://seqco.de/i:49710

**Description of *Acaudatibacter aquilonius* sp. nov.**

*Acaudatibacter aquilonius* (a.qui.lo'ni.us. L. masc. adj. *aquilonius*, northern, northerly, referring to the recovery of genomes of the organism from northern freshwater bodies).

Twelve metagenome-assembled genomes representing this species were assembled from sequence data obtained from samples taken from lakes Björntjärnen (Sweden), Alinen Mustajärvi (Finland), Keskinen Rajajärvi (Finland), and Valkea Kotinen (Finland). Completeness estimates for genomes, as determined by CheckM (v1.1.3; 'lineage_wf'), are 69.50–97.53%, with 1.62–4.21% estimated contamination. Genome assemblies range between 3.06–4.56 Mbp in size, comprising 100–973 contigs, with a G+C content of 67.37%–67.58%. Estimated complete genome sizes from CheckM range between 4.40–4.68 Mbp. ANI and AAI values between these genomes are 97.8%–100% and 99.3–100%, respectively, while such pairwise comparisons to closely related taxa are below 81.0% and 78.8%, respectively. Phylogenomic analysis of 72 concatenated conserved alphaproteobacterial single-copy genes places this species in the genus *Acaudatibacter*, in the family *Caulobacteraceae*. The species corresponds to GTDB taxonomy (R220) species "Palsa-881 sp903872075".

Genomes lack multiple genes for flagellar motility, chemotaxis, holdfast adhesin production, and for the caulobacterial obligate dimorphic cell developmental program. Genomes contain genes for type IV tight-adhesion pili; for carotenoid pigment production; for complete biosynthesis pathways of all standard amino acids; for aerobic respiration using cytochrome *c* oxidases cytochrome $aa_3$ (*coxABC*) and cytochrome $cbb_3$ (*ccoNOPQ*); for thiosulfate oxidation using thiosulfate dehydrogenase TsdA; for biosynthesis and degradation of polyphosphate and polyhydroxybutyrate; and for high-affinity PstABCS phosphate and PhnDEC phosphonate transporters. In addition, they have genetic potential for photoautotrophy, containing genes for bacteriochlorophyll synthesis, type II anoxygenic photosynthesis using a light-harvesting II (LH2) complex and a reaction center–light-harvesting I supercomplex (RC–LH1), as well as carbon fixation using the Calvin-Benson-Bassham (CBB) cycle, including the accessory genes for red-type RuBisCO activase (*cbbX*) and XuBP phosphatase (*cbbY*). The species has been detected in both oxic and anoxic strata of stratified freshwater bodies in Finland and Sweden.

Likely a psychrophilic/mesophilic facultative anaerobe, based on its genetic repertoire and environmental distribution.

The proposed nomenclatural type for the species is the genome Umea_bin-04329$^{Ts}$, available under the NCBI WGS assembly accession number GCA_903872075.1$^{Ts}$ (BioProject ID PRJEB38681), recovered as a metagenome coassembly from samples taken in autumn of 2018 from the stratified lakes Björntjärnen (lat. 64.12, long. 18.78) and Nästjärnen (lat. 64.15, long. 18.80), both in Umeå, Sweden (metagenomes ERS4600559–565 and ERS4600568–570). It comprises 100 contigs with a total of 4,361,582 bp, and has an estimated completeness of 95.32% and contamination of 1.62%.

SeqCode URL: https://seqco.de/i:49709

**Description of *Acaudatibacter boreus* sp. nov.**

*Acaudatibacter boreus* (bo're.us. L. masc. adj. *boreus*, northern, referring to the recovery of genomes of the organism from northern freshwater bodies).

Four metagenome-assembled genomes representing this species were assembled from sequence data obtained from samples taken from anoxic strata of a lake in Kiruna (Sweden), referred to as Ki1. Completeness estimates for genomes, as determined by CheckM (v1.1.3; 'lineage_wf'), are 93.45%–94.46%, with 2.03–3.83% estimated contamination. Genome assemblies range between 4.23 Mbp–4.36 Mbp, comprising 576–699 contigs, with a G+C content of 67.46%–67.60%. Estimated complete genome sizes from CheckM range between 4.52–4.62 Mbp. ANI and AAI values between these genomes are 99.2%–100% and 99.6–100%, respectively, while such pairwise comparisons to closely related taxa are below 80.9% and 77.9%, respectively. Phylogenomic analysis of 72 conserved alphaproteobacterial single-copy genes places this species in the genus *Acaudatibacter*, in the family *Caulobacteraceae*. The species corresponds to GTDB taxonomy (R220) species "Palsa-881 sp903870555".

Genomes lack multiple genes for flagellar motility, chemotaxis, holdfast adhesin production, and for the caulobacterial obligate dimorphic cell developmental program. Genomes contain genes for type IV tight-adhesion pili; for carotenoid pigment production; for complete biosynthesis pathways of all standard amino acids; for aerobic respiration using cytochrome *c* oxidases cytochrome $aa_3$ (*coxABC*) and cytochrome $cbb_3$ (*ccoNOPQ*), and ubiquinol oxidase cytochrome *bd* (*cydABX*); for thiosulfate oxidation using the thiosulfate dehydrogenase TsdA; for sulfide oxidation using the Sqr sulfide:quinone oxidoreductase; for biosynthesis and degradation of polyphosphate, polyhydroxybutyrate, and starch/glycogen; and for the high-affinity PstABCS phosphate, PhnDEC phosphonate, and SsuABC sulfonate transporters. In addition, they have partial genetic potential for photoautotrophy, containing genes for type II anoxygenic photosynthesis using a light-harvesting II (LH2) complex and a reaction center–light-harvesting I supercomplex (RC–LH1), as well as partial genetic potential for carbon fixation using the Calvin-Benson-Bassham (CBB) cycle. The species has been detected in both oxic and anoxic strata of stratified freshwater bodies in Canada, Finland, and Sweden. Likely a psychrophilic/mesophilic facultative anaerobe, based on its genetic repertoire and environmental distribution.

The proposed nomenclatural type for the species is the genome Ki1-2-2m_bin-386$^{Ts}$, available under the NCBI WGS assembly accession number GCA_903870555.1$^{Ts}$ (BioProject ID PRJEB38681), recovered 24 July 2018 from a stratified lake in Kiruna, Sweden (lat. 67.93, long. 20.36; from the metagenome ERS4600419). It comprises 692 contigs with a total of 4,349,168 bp, and has an estimated completeness of 94.45% and contamination of 2.03%.

SeqCode URL: https://seqco.de/i:49711

**Description of *Acaudatibacter lapponiensis* sp. nov.**

*Acaudatibacter lapponiensis* (lap.po.ni.en'sis. N.L. masc. adj. *lapponiensis*, pertaining to Lapponia, the Latin name for Lapland, the geographical region from which genomes of the organism were recovered).

Four metagenome-assembled genomes representing this species were assembled from sequence data obtained from samples taken from a lake in Kiruna (Sweden), referred to as Ki2, Swedish lake code: 754378-169136. Completeness estimates for genomes, as determined by CheckM (v1.1.3; 'lineage_wf'), are 94.86%–96.97%, with 1.94%–4.47% estimated contamination. Genomes assemblies range between 4.57–4.64 Mbp, comprising 279–357 contigs, with a G+C content of 68.83%–68.84%. Estimated complete genome sizes from CheckM range between 4.73–4.81 bp. ANI and AAI values between these genomes are 99.9%–100% and 99.9–100%, respectively, while such pairwise comparisons to closely related taxa are below 81.0% and 78.8%, respectively. Phylogenomic analysis of 72 conserved alphaproteobacterial single-copy genes places this species in the genus *Acaudatibacter*, in the family *Caulobacteraceae*. The species corresponds to GTDB taxonomy (R220) species "Palsa-881 sp903923135".

Genomes lack multiple genes for flagellar motility, chemotaxis, holdfast adhesin production, and for the caulobacterial obligate dimorphic cell developmental program. Genomes contain genes for type IV tight-adhesion pili; for carotenoid pigment production; for complete biosynthesis pathways of all standard amino acids; for aerobic respiration using cytochrome *c* oxidases cytochrome $aa_3$ (*coxABC*) and cytochrome $cbb_3$ (*ccoNOPQ*), and for ubiquinol oxidase cytochrome *bd* (*cydA*; partial genetic potential); for biosynthesis and degradation of polyphosphate, polyhydroxybutyrate, and starch/glycogen; for the high-affinity PstABCS phosphate, PhnDEC phosphonate, and SsuABC sulfonate transporters. In addition, they have genetic potential for photoautotrophy, containing genes for type II anoxygenic photosynthesis using a light-harvesting II (LH2) complex and a reaction center–light-harvesting I supercomplex (RC–LH1), as well as carbon fixation using the Calvin-Benson-Bassham (CBB) cycle, including the accessory genes for red-type RuBisCO activase (*cbbX*) and XuBP phosphatase (*cbbY*). The species has been detected in both oxic and anoxic strata of stratified freshwater bodies in Finland and Sweden. Likely a psychrophilic/mesophilic facultative anaerobe, based on its genetic repertoire and environmental distribution.

The proposed nomenclatural type for the species is the genome Kiruna2_bin-0871$^{Ts}$, available under the NCBI WGS assembly accession number GCA_903923135.1$^{Ts}$ (BioProject ID PRJEB38681), recovered as a coassembly from samples taken 27 July 2018 from a stratified lake in Kiruna, Sweden (lat. 67.92, long. 20.37; from the metagenomes ERS4600421–424). It comprises 279 contigs with a total of 4,636,946 bp, and has an estimated completeness of 96.47% and contamination of 2.20%.

SeqCode URL: https://seqco.de/i:49712

### SUPPLEMENTARY FIGURES

**Figure S1**. Maximum-likelihood (ML) species phylogeny of 72 concatenated and manually curated single-copy marker genes[25] for all 347 *Caulobacterales* species genome representatives, the model organism *C. crescentus* (*C. vibrioides*) CB15, and five outgroup *Alphaproteobacteria* for rooting. Inferred using IQ-TREE[30] with the LG+C60+F+R model of evolution and 100 non-parametric bootstraps (alignment length of 25448 amino acids). Family and genus assignment follows GTDB taxonomy (R207)[31], with minor modifications according to the taxonomic proposals in **Supplementary Discussions 1–2**. Same tree as shown in **Fig. 1a**, but with all species labeled.

**Figure S2**. ML species phylogeny of 117 alphaproteobacterial concatenated single-copy marker genes extracted using GToTree[24], for all 347 *Caulobacterales* species genome representatives, the model organism *C. crescentus* (*C. vibrioides*) CB15, and five outgroup *Alphaproteobacteria* for rooting. Inferred using IQ-TREE[30] with the LG+C60+F+G model of evolution and 1000 ultrafast bootstraps (alignment length of 21766 amino acids). Family and genus assignment follows GTDB taxonomy (R207)[31], with minor modifications according to the taxonomic proposals of **Supplementary Discussions 1–2**.

**Figure S3**. Representative micrographs used for schematic drawings in **Figs. 1a** and **4d**, of cells sampled at the specified growth conditions. Scale bar: 5 μm.

**Figure S4**. **(a)** Species phylogeny shown in **Fig. 1a**. Numbers represent non-parametric bootstraps and the scale bar indicates number of substitutions per site. (**b–j**) Expanded view of the presence and absence of genes presented in **Fig. 2b**, showing genes involved in (b) chemotaxis, (c) flagellum, (d) cell cycle and developmental genes, (e) type IV adhesive pilus (T4P), (f) holdfast, (g) crescentin, (h) S-layer, (i) prostheca, and (j) cell division, among *Caulobacterales* genomes. Gene orthologs were identified using KEGG ortholog (KO) annotations from eggNOG-mapper[32] v2.1.5 (dark gray) or through the reciprocal best blast hit (RBH) algorithm using the *C. crescentus* CB15 proteome (blue). For RBH results, the corresponding loci in the *C. crescentus* CB15 (CC numbers) and *C. crescentus* NA1000 (CCNA numbers) are shown alongside the gene name. Descriptions come from the KO annotation or from the *C. crescentus* NA1000 genome annotation. Numbers show KO copy numbers > 1. The full dataset is found in **Supplementary Data S4**.

**Figure S5**. Overview of metadata for the subset of *Acaudatibacter* gen. nov. genomes included in Rodríguez-Gijón *et al.*[33] here referred to as the "extended dataset". **(a)** Genome attributes for the 34 *Acaudatibacter* metagenome-assembled genomes (MAGs). Taxon names are written in bold font and MAG IDs in light font. Five-digit species cluster names match the dataset of Rodríguez-Gijón *et al.*[33] *: MAGs used as species genome representatives in our "core dataset". **(b)** Metagenome sampling site metadata based on NCBI BioSample information, and from Nayfach *et al.*[34] for assemblies 3300015024_21 and 3300025574_7, presented as done in **Fig. 1b**. See panel (c) for environment color codes. Metadata are presented in the right-hand table for MAGs obtained from individual freshwater metagenomes[35] for which water chemistry measurements are available. **(c)** Summary of the phylogenetic relationship, genetic potential, and habitats of the twelve *Acaudatibacter* species clusters, alongside *C. crescentus* and *P. immobile*. **: putative dimorphic or monomorphic lifecycles is inferred from the presence or large absence of developmental regulators of the dimorphic lifecycle, respectively (**Fig. 2b**, **Extended Data Fig. 3**), and the reproductive symmetry of *P. immobile* (**Fig. 3**). Circles represent genetic potential shown in **Extended Data Fig. 3** and **Supplementary Figs. S9**, **S11–S12**. Full phylogeny is presented in **Extended Data Fig. 4**.

**Figure S6**. Genomes of putatively monomorphic *Caulobacteraceae* species encode canonical c-di-GMP enzymes. **(a)** The number of GGDEF (c-di-GMP biosynthesis) and/or EAL domain (c-di-GMP degradation) proteins per genome, shown for *Caulobacteraceae* genera. Putatively monomorphic species are highlighted in color alongside the model strain *C. crescentus* CB15. Black vertical lines depict mean counts for each genus. Protein domains were identified using InterProScan[36]. **(b–c)** Proteins containing EAL (b) or GGDEF (c) domains aligned using Clustal Omega[20]. Most proteins show canonical, non-degenerate EAL (Glu-Ala-Leu) or GGDEF (Gly-Gly-Asp/Glu-Glu-Phe) motifs, with most GGDEF domain proteins additionally containing the allosteric I-site motif (Arg-x-x-Asp)—key amino acid residues involved in the catalysis of c-di-GMP degradation or biosynthesis[37]. Non-functional c-di-GMP biosynthesis enzymes typically contain degenerate GGDEF motifs[37].

**Figure S7**. **(a)** ML phylogeny of crescentin (CreS) homologs. The phylogeny was inferred using IQ-TREE[30] with the Q.pfam+C60+R9 model of evolution and 1000 ultrafast bootstraps (356 sequences with 902 amino acid positions), and was rooted with the major *Beta-/Gammaproteobacteria* clade. Same tree as shown in **Fig. 4c**. **(b)** Pfam protein domains identified for each protein included in (a). Proteins without predicted domains are not shown. For collapsed clades in **Fig. 4c**, the major taxonomic group is specified alongside the numbers of sequences per clade shown in parentheses. The scale bar indicates the number of substitutions per site.

**Figure S8**. ML phylogeny including only the clade from **Fig. 4c** and **Supplementary Fig. S7** where proteins with predicted crescentin domains (PF19220) are found. The phylogeny was inferred using IQ-TREE[30] with the JTT+C60+R5 model of evolution and 1000 ultrafast bootstraps (135 sequences with 407 amino acid alignment positions), and was rooted between *Hyphomicrobiales* and *Caulobacteraceae*. The scale bar indicates number of substitutions per site. *Hyphomicrobiales* families and *Caulobacteraceae* genera are indicated. Same tree as shown in **Fig. 4d**.

**Figure S9**. **(a)** Species phylogeny shown in **Fig. 1a**. Numbers represent non-parametric bootstraps and the scale bar indicates number of substitutions per site. Genomes containing phototrophy genes are marked with red circles. **(b)** Meta-analysis of colony pigments across *Caulobacterales* species. See **Supplementary Data S6** for colony descriptor words and references. Asterisks: *Caulobacter* isolates ErkDOM-C and ErkDOME pigment descriptions derive from this work. **(c–i)** Expanded view of the presence and absence of genes presented in **Fig. 5b**, showing genes involved in (c) carotenoid biosynthesis, (d) bacteriochlorophyll biosynthesis, (e) bacteriochlorophyll transport, (f) light-harvesting complex II (LH2), (g) reaction center–light-harvesting complex I (RC–LH1), (h) $CO_2$ fixation using the CBB cycle, and (i) aerobic respiration among *Caulobacterales* genomes. KEGG ortholog (KO) gene ortholog were annotated using either eggNOG-mapper[32] (emapper; v2.1.5) or GhostKOALA[38] (v2.2). Numbers show KO copy numbers > 1. Abbreviations: cytochrome (cyt.). Full dataset is found in **Supplementary Data S4**. **(j)** Schematic representation of the highly branched electron transport chain of *C. crescentus* CB15, which includes two high-affinity terminal oxidases operating under low-oxygen concentrations (cytochromes *bd* and $bb_3$) and two low-affinity terminal oxidases operating under high-oxygen concentrations (cytochromes $bo_3$ and $aa_3$)[39].

**Figure S10**. Gene synteny of *Caulobacterales* photosynthesis gene clusters (PGCs). **(a)** Cladogram representation of the PufM phylogeny shown in **Fig. 5e** and **Supplementary Fig. S13**, pruned to only include relevant species. Ultrafast bootstrap proportions (ufBPs) are indicated with circles. Single asterisks highlight species absent from the PufM phylogeny; their vertical placement here is based instead on the BchY phylogeny of **Supplementary Fig. S15**. Double asterisks highlight the taxon "REEB506" sp. REEB506, for which its PufM ortholog was placed without support outside of the main *Caulobacterales* clade (**Supplementary Fig. S13**); its vertical placement here close to the taxa "UBA4763" and "REEB509" is based instead of the BchY phylogeny of **Supplementary Fig. S15**. **(b)** Synteny of *Caulobacterales* phototrophy genes. Species have been sorted vertically following the topology of panel (a). Genes are color-coded and schematically labeled with letters corresponding to their gene names, and orthologs are linked with gray boxes. Contigs are marked with "c" followed by the contig number and distances between loci on the same contig are shown with black boxes. When present and fully enclosed within 5 kb of a phototrophy gene, additional flanking genes are shown to highlight broader genetic context. Uncharacterized genes coding for PF02655 (ATP-grasp domain) and cytochrome P450 proteins, as well as for DUF3422 (domain of unknown function 3422) proteins, were tentatively labeled as carotenoid- and bacteriochlorophyll-associated, respectively, based on their colocalization patterns.

**Figure S11**. Presence and absence of genes, in *Acaudatibacter* gen. nov. "extended dataset" genomes sourced from Rodríguez-Gijón *et al.*[33], for pigment production, phototrophy, carbon fixation, and aerobic respiration. **(a–g)** Presence and absence of KEGG orthologs (KOs) annotated using either eggNOG-mapper[32] (emapper; v2.1.12) or GhostKOALA[38] (v3.0), involved in (a) carotenoid biosynthesis, (b) bacteriochlorophyll biosynthesis, (c) bacteriochlorophyll transport, (d) light-harvesting complex II (LH2), (e) reaction center–light-harvesting complex I (RC–LH1), (f) $CO_2$ fixation via the CBB cycle, and (g) aerobic respiration among *Caulobacterales* genomes. Numbers show KO copy numbers $> 1$. Abbreviations: cytochrome (cyt.). The full dataset can be found in **Supplementary Data S4**.

**Figure S12**. Estimated completeness of carbon fixation pathways among *Caulobacterales* genomes based on KEGG ortholog (KO) annotations from eggNOG-mapper[32] (v2.1.5) and associated KEGG modules. **(a)** Species phylogeny shown in **Fig. 1a**. Numbers represent non-parametric bootstraps and the scale bar indicates number of substitutions per site. Genomes containing phototrophic potential are marked with red circles for *Caulobacterales* and with dark blue circles for outgroup *Alphaproteobacteria*. Genomes with complete genetic potential for the CBB cycle are marked with orange circles. **(b)** Calvin-Benson-Bassham (CBB) cycle—KEGG module M00165. *Left panel*: Completeness of the CBB cycle steps. *Right panel*: Copy number of individual CBB cycle KOs. **(c)** Completeness of the reductive citrate cycle steps—KEGG module M00173. **(d)** Completeness of the 3-hydroxy-propionate bicycle steps—KEGG module M00376.

**Figure S13**. ML phylogeny of photosynthetic reaction center protein PufM, inferred using IQ-TREE[30] with the WAG+C60+R9 model of evolution and 1000 ultrafast bootstraps (278 sequences with 295 amino acid alignment positions). The tree was rooted at the major *Chromatiales* (*Gammaproteobacteria*) clade. Taxonomic groups are color-coded as outlined in the color legend. The scale bar indicates number of substitutions per site. Same tree as shown in **Fig. 5e**; gradients highlight collapsed clades in **Fig. 5e**.

**Figure S14**. ML phylogeny of photosynthetic reaction center protein PufM, inferred as in **Supplementary Fig. S13**, but with less stringent alignment trimming (278 sequences with 324 amino acid positions). The tree was rooted at the major *Chromatiales* (*Gammaproteobacteria*) clade. Taxonomic groups are color-coded as outlined in the color legend. The scale bar indicates number of substitutions per site. Same tree as shown in **Extended Data Fig. 7a**; gradients highlight collapsed clades in **Extended Data Fig. 7a**.

**Figure S15**. ML phylogeny of chlorophyllide *a* reductase subunit BchY, inferred using IQ-TREE[30] with the WAG+C60+R10 model of evolution and 1000 ultrafast bootstraps (352 sequences with 417 amino acid alignment positions). The tree was rooted at the major *Chromatiales* (*Gammaproteobacteria*) clade. Taxonomic groups are color-coded as outlined in the color legend. The scale bar indicates number of substitutions per site. Same tree as shown in **Extended Data Fig. 7b**; gradients highlight collapsed clades in **Extended Data Fig. 7b**.

**Figure S16**. ML phylogeny of chlorophyllide *a* reductase subunit BchY, inferred as in **Supplementary Fig. S13**, but with less stringent alignment trimming (352 sequences with 509 amino acid alignment positions). The tree was rooted at the major *Chromatiales* (*Gammaproteobacteria*) clade. Taxonomic groups are color-coded as outlined in the color legend. The scale bar indicates number of substitutions per site. Same tree as shown in **Extended Data Fig. 7c**; gradients highlight collapsed clades in **Extended Data Fig. 7c**.

**Figure S17**. Geographical distribution of freshwater *Acaudatibacter* species. **(a)** Competitive mapping results of *Caulobacteraceae* species clusters (genomes of > 50% completeness and < 5% contamination, clustered at > 95% ANI) against 636 freshwater metagenomes, from Rodríguez-Gijón *et al.*[33] Out of the 325 *Caulobacteraceae* species clusters investigated, 190 were present in at least one freshwater metagenome (red circles). Dark blue circles represent metagenomes to which none of the 325 *Caulobacteraceae* species clusters mapped. Note that some locations have multiple geographically proximal freshwater bodies and/or multiple metagenomes per freshwater body, resulting in overlapping datapoints. **(b)** Same as in (a), but mapping only the subset of species clusters (n = 12) belonging to the genus *Acaudatibacter* gen. nov. (GTDB taxon "g__Palsa-881"). The five *Acaudatibacter* species clusters that mapped to freshwater metagenomes are indicated. **(c)** Total sum of relative abundances across sampled depths per water body, of the twelve *Acaudatibacter* species clusters among metagenomes from 41 stratified freshwater bodies represented in the dataset of Buck *et al.*[35] (see **Fig. 6c** and **Extended Data Fig. 9**), as a proxy for species cluster distribution. Genomes of the species clusters labeled in black did not map to the dataset. Among the twelve *Acaudatibacter* species clusters investigated, five are represented in our core dataset of 347 *Caulobacterales* species genome representatives, as indicated.

**Figure S18**. Presence (dark gray) and absence (light gray) of flagellar and type IV tight-adhesion pili (T4P) KEGG orthologs (KOs) predicted using eggNOG-mapper[32] (v2.1.12) among curved *creS*-encoding *Hyphomicrobiales* species. Numbers show KO copy numbers above one.

### SUPPLEMENTARY TABLES

**Table S1**. List of published micrographs used as reference for drawing schematic illustrations in **Figs. 1a** and **4d**.

| | Micrograph used as guide for drawing | | |
|---|---|---|---|
| **Species** | **Figure/file in orig. publ.** | **Reference** | **Modifications made to cell outline** |
| *P. haematophilum* | Figure 2a | 40 | |
| *P. glaciei* | Figure 2 | 41 | |
| *P. parvum* | Figure S1 | 42 | |
| *C. rhizosphaerae* | Figure S1* | 43 | *Figs. S1a and S1b of Sun *et al.*[43] were combined; prostheca based on Fig. S1b added to predivisional cell outline based on Fig. S1a. |
| *Po. montana* comb. nov. (*P. montanum*) | Figure 2 | 5 | |
| *B. vesicularis* | Figure 2 | 44 | |
| *B. mediterranea* | Figure 2 | 44 | |
| *B. nasdae* | Figure 1 | 44 | |
| *B. terrae* | Figure 2 | 44 | |
| *B. pondensis* | Figure 4a | 45 | |
| *B. basaltis* | Figure 2 | 44 | Prostheca based on non-predivisional cell was added to cell outline based on predivisional cell. |
| *B. denitrificans* | Figure 2 | 46 | |
| *B. aveniformis* | Figure 1 | 44 | |
| *A. tiandongensis* | Figure S1 | 47 | |
| *A. benevestitus* | Figure 1b | 48 | |
| *A. taihuensis* | Figure 1 | 49 | |
| *Hyphomonas neptunia* | Figure 6c | 50 | |
| *Hyphomonas adhaerens* | Figure 1 | 51 | |
| *Henriciella barbarensis* | Figure 2 | 52 | |
| *Ponticaulis koreensis* | Figure 2 | 53 | |
| *Hirschia baltica* | Figure 1 | 54 | |
| *Vitreimonas silvestris* comb. nov. (*Terricaulis silvestris*) | Figure S1B | 4 | |
| *Vitreimonas flagellata* | Figure S1 | 3 | |
| *Pseudaquidulcibacter paucihalophilus* | Figure 1b | 2 | |
| *"Maricaulis alexandrii"* | Figure 1d | 55 | |
| *Hyphomicrobium indicum* | Figure 2a | 56 | |
| *Glycocaulis alkaliphilus* | Figure 1a | 57 | |
| *Marinicauda pacifica* | Figure S2a | 58 | |
| *Oceanicaulis alexandrii* | Figure 3a | 17 | |
| *Oceanicaulis satelles* comb. nov. (*Alkalicaulis satelles*) | Figure 1a | 6 | |
| *Woodsholea maritima* | Figure 1a | 59 | |
| *Robiginitomaculum antarcticum* | Figure 2a | 60 | Prostheca-like protrusion seen on non-predivisional cell was added to cell outline based on predivisional cell. |
| *Parvularcula bermudensis* | Figure 1a | 61 | |
| *Aquisalinus flavus* | Figure S1 | 62 | |
| *Amphiplicatus metrithermophilus* | Figure S1a | 63 | |
| *Methylocystis parva* | Figure BXII.α.153 | 64 | |
| *Methylocystis* sp. Rockwell | Figure 2a | 65 | |
| *Roseiarcus fermentans* | Figure 1a | 66 | |

**Table S2**. List of strains used in this work. Abbreviations: genta, gentamicin; DSMZ, German Collection of Microorganisms and Cell Cultures; CCUG, Culture Collection University of Gothenburg.

| Species | # | Strain | Background | Genotype | Medium | Source | Ref. |
|---|---|---|---|---|---|---|---|
| *Aquidulcibacter paucihalophilus* | KJ1143 | DSM 109892$^T$ [TH1-2$^T$] | | | PYE | DSMZ | |
| *Asticcacaulis biprosthecius* | KJ1167 | DSM 4723$^T$ [C19$^T$] | | | PYE | DSMZ | |
| *Asticcacaulis excentricus* | KJ1168 | DSM 4724$^T$ [CB48$^T$] | | | PYE | DSMZ | |
| *Brevundimonas aurantiaca* | KJ1149 | CCUG 45020$^T$ [CB-R$^T$] | | | PYE | CCUG | |
| *Brevundimonas bacteroides* | KJ998 | DSM4726$^T$ [CB7$^T$] | | | PYE | DSMZ | |
| *Brevundimonas diminuta* | KJ1144 | DSM 7234$^T$ [Pickett K-248$^T$] | | | PYE | DSMZ | |
| *Brevundimonas goettingensis* | KJ1145 | DSM 112305$^T$ [LVF2$^T$] | | | PYE | DSMZ | |
| *Brevundimonas lenta* | KJ1146 | DSM 23960$^T$ [DS-18$^T$] | | | PYE | DSMZ | |
| *Brevundimonas subvibrioides* | KJ1148 | DSM 4735$^T$ [CB81$^T$] | | | PYE | DSMZ | |
| *Brevundimonas variabilis* | KJ997 | DSM4737$^T$ [CB17$^T$] | | | PYE | DSMZ | |
| *Caulobacter crescentus* (*C. vibrioides*) | KJ883 | CB15 | | | PYE | S. Crosson | |
| *Caulobacter crescentus* (*C. vibrioides*) | KJ1 | NA1000 | | | PYE | M. Laub | [67] |
| *Caulobacter crescentus* (*C. vibrioides*) | KJ1183 | | NA1000 | pBXMCS-4 (EV) | PYE + genta | This work | |
| *Caulobacter crescentus* (*C. vibrioides*) | KJ1184 | | NA1000 | pBXMCS-4-$P_{xyl}$-*creS* | PYE + genta | This work | |
| *Caulobacter crescentus* (*C. vibrioides*) | KJ1185 | | NA1000 | pBXMCS-4-$P_{xyl}$-*creS*$_{Ch.reniformis}$ | PYE + genta | This work | |
| *Caulobacter crescentus* (*C. vibrioides*) | KJ1179 | CJW1208 (LS3812) | NA1000 | Δ*creS* | PYE | C. Jacobs-Wagner | [68] |
| *Caulobacter crescentus* (*C. vibrioides*) | KJ1186 | | CJW1208 | Δ*creS* pBXMCS-4 (EV) | PYE + genta | This work | |
| *Caulobacter crescentus* (*C. vibrioides*) | KJ1187 | | CJW1208 | Δ*creS* pBXMCS-4-$P_{xyl}$-*creS* | PYE + genta | This work | |
| *Caulobacter crescentus* (*C. vibrioides*) | KJ1188 | | CJW1208 | Δ*creS* pBXMCS-4-$P_{xyl}$-*creS*$_{Ch.reniformis}$ | PYE + genta | This work | |
| *Caulobacter flavus* | KJ1169 | DSM 29968$^T$ [RHGG3$^T$] | | | PYE | DSMZ | |
| *Caulobacter henricii* | KJ1151 | CCUG 49339$^T$ [CB4$^T$] | | | PYE | CCUG | |
| *Caulobacter mirabilis* | KJ1170 | DSM 21795$^T$ [FWC38$^T$] | | | PYE | DSMZ | |
| *Caulobacter segnis* | KJ1147 | DSM 7131$^T$ [TK0059$^T$] | | | PYE | DSMZ | |
| *Caulobacteraceae* sp. PMMR1 | KJ1174 | DSM 26776 [PMMR1] | | | R2A pH 6.0 | DSMZ | |
| *Chelatococcus reniformis* | KJ1171 | DSM 105737$^T$ [B2974$^T$] | | | PYE | DSMZ | |
| *Escherichia coli* | KJ665 | MG1655 | K12 | $F^-$ $\lambda^-$ *ilvG*$^-$ *rfb*-50 *rph*-1 | LB | M. Laub | |
| *Escherichia coli* | N/A | DH5α | | [general cloning strain] | LB | Invitrogen | |
| *Escherichia coli* | N/A | | TOP10 | pBXMCS-4 (EV) | LB + genta | M. Thanbichler | [69] |
| *Escherichia coli* | KJ1181 | | DH5α | pBXMCS-4-$P_{xyl}$-*creS* | LB + genta | This work | |
| *Escherichia coli* | KJ1182 | | DH5α | pBXMCS-4-$P_{xyl}$-*creS*$_{Ch.reniformis}$ | LB + genta | This work | |
| *Phenylobacterium deserti* | KJ1172 | DSM 103871$^T$ [YIM 73061$^T$] | | | R2A | DSMZ | |
| *Phenylobacterium immobile* | KJ1173 | DSM 1986$^T$ [E$^T$] | | | R2A | DSMZ | |

**Table S3**. List of oligonucleotides used in this work.

| Name | Sequence (5′–3′) |
|---|---|
| OJH25 | atggtcgtctccccaaaactc |
| OJH26 | ctgcagcccgggggatccactag |
| OJH27 | gctcgagttttggggagacgaccatatgagactgctgtcgaaga |
| OJH28 | aactagtggatcccccgggctgcagttaggcgctcgcggccacg |
| OJH29 | gctcgagttttggggagacgaccatatgatcggtattggacagc |
| OJH30 | aactagtggatcccccgggctgcagttattcggccgcggtattg |

#### SUPPLEMENTARY VIDEOS

**Supplementary Video 1**. Timelapse microscopy of *P. immobile* $E^T$ at 30°C on R2A 1% agarose, in 5-minute imaging intervals. Playback speed: 18 frames/s (1.5 h/s). Scale bars: 1 µm. File format: .avi. **(a)** The cell shown in **Fig**. **3a**. **(b)** Three additional cells.

**Supplementary Video 2**. Timelapse microscopy of *E. coli* MG1655 at 37°C on LB 1% agarose, in 20-second imaging intervals. Playback speed: 30 frames/s (15 min/s). Scale bars: 1 µm. File format: .avi. **(a)** The cell shown in **Fig**. **3a**. **(b)** Three additional cells.

**Supplementary Video 3**. Timelapse microscopy of *C. crescentus* CB15 at 30°C on PYE 1% agarose, in 30-second imaging intervals. Playback speed: 30 frames/s (15 min/s). Scale bars: 1 µm. File format: .avi. **(a)** The cell shown in **Fig**. **3a**. **(b)** Six additional cells.

#### SUPPLEMENTARY DATA

**Data S1 | Genome overview.** Overview of genomes used in this work. **(a)** The "core dataset" of *Caulobacterales* genomes. Columns A–C, genome name, accession, and taxon name; Columns D–G, information on genome dereplication and selection of species representatives; Column H, GTDB taxonomy (release R207); Columns I–R, genome statistics, including assembly size, estimated genome size, N50, number of contigs, G+C content, and estimates of completeness and contamination from both the CheckM[70] methods 'taxonomy_wf' (used for genome selection) and 'lineage_wf' (used for estimated genome size calculation); Columns S–U, additional information. **(b)** The "extended dataset" of *Acaudatibacter* gen. nov. (GTDB taxon "g__Palsa-881") genomes sourced from Rodríguez-Gijón *et al.*[33] Column A, genome name; Column B, references for assemblies[33-35, 71-74]; Column C–D, species clustering information; Column E, whether the genome is also included as a species genome representative (SGR) in the "core dataset"; Column F, type of genome (MAG, metagenome-assembled genome); Column G, GTDB taxonomy (release R207); Columns H–N, genome statistics, including assembly size, estimated genome size, number of contigs, number of scaffolds, G+C content, and estimates of completeness and contamination from the CheckM method 'lineage_wf'.

**Data S2 | Environmental metadata.** Meta analysis of genome sampling environment metadata. **(a)** Explanation of the table layout and content of the metadata. **(b)** Manually collected and curated metadata from NCBI BioSample (www.ncbi.nlm.nih.gov/biosample/) and BioProject (www.ncbi.nlm.nih.gov/bioproject/) pages, as well as JGI Gold (gold.jgi.doe.gov/), and listed publications when necessary. Listed literature references: [2, 6, 7, 9-12, 17, 21, 22, 28, 40, 43, 49, 51, 52, 55, 59, 61-63, 75-116].

**Data S3 | IMNGS environmental data.** Compiled IMNGS[117] 'Taxonomy' job results for the query: "Bacteria/Proteobacteria/Alphaproteobacteria/Caulobacterales/Caulobacteraceae".

**Data S4 | Selected gene presence/absence data.** Gene presence and absence data presented in figures and supplementary figures of the article (**Figs. 2b**, **4b**, and **5b**, **Extended Data Figs. 2** and **3**, as well as **Supplementary Figs. S4**, **S9**, and **S11)**. Includes genes for chemotaxis, flagellar motility, cell cycle and development, type IV pilus, holdfast synthesis, crescentin, S-layer, prostheca, cell division, carotenoid synthesis, photosynthesis, carbon fixation, aerobic respiration, secretion systems, and sulfonate transport. Includes both *Caulobacterales* genomes of the "core dataset", and *Acaudatibacter* genomes of the "extended dataset". For gene orthologs, attributes are separated by "@" in the following order: (1) gene category, (2) annotation tool, (3) KEGG KO or *C. crescentus* locus IDs, (4) gene name, (5) EC number, (6) gene annotation, (7) manually curated gene name. For RBH results, the attributes #4 and #7 were taken from the *C. crescentus* NA1000 GCF_000022005.1 assembly, since it is better annotated, but they all agree well with the *C. crescentus* CB15 GCF_000006905.1 assembly. For genomes, attributes are separated by "@" in the following order: (1) dataset [either the "core" dataset of *Caulobacterales* species genome representatives or the "extended" dataset of additional *Acaudatibacter*/"Palsa-881" genomes], (2) assembly ID, (3) family, (4) genus, (5) species, (6) taxon name for the genome assembly [only for the "extended" dataset]. KOs of the "core" dataset were annotated using eggNOG-mapper[32] (emapper) v2.1.5 or GhostKOALA[38] v2.3 and KOs of the "extended" dataset were annotated using eggNOG-mapper (emapper) v2.1.12 or GhostKOALA v3.0.

**Data S5 | Uncharacterized putative flagellar/developmental genes.** Identification of putative flagellar motility and development factors, based on their absence from non-flagellated *Acaudatibacter* gen. nov. and *Phenylobacterium* species. Presence/absence is based on the reciprocal best blast hit (RBH) algorithm. The 100 genes missing from non-flagellated lineages, sorted by their conservation in the *Acaudatibacter–Caulobacter–Phenylobacterium* (ACP) clade. For gene orthologs, attributes are separated by "@" in the following order: (1) gene category, (2) *C. crescentus* CB15 protein accession, (3) CB15 new locus ID, (4) CB15 old locus ID, (5) CB15 gene name, (6) CB15 gene annotation, (7) *C. crescentus* NA1000 protein accession, (8) NA1000 locus ID, (9) NA1000 gene name, (10) NA1000 gene annotation, (11) total number of species genome representatives having the gene within the ACP clade [basis for sorting]. For genomes, attributes are separated by "@" as presented in **Supplementary Data S4**.

**Data S6 | Colony pigments.** Meta analysis of the colony pigment description terms used in the literature for *Caulobacterales* isolates included in our dataset, if available. Listed literature references: [1-4, 6, 7, 9, 17, 21, 22, 28, 29, 40-43, 45-49, 51-60, 62, 63, 77, 78, 81, 83, 85, 88-90, 92-97, 99, 100, 106, 107, 109-111, 113, 115, 118-144].

**Data S7 | Overview of eggNOG-mapper-predicted NOGs.** Overview of eggNOG-mapper[32] (emapper) annotations of non-supervised orthologous groups (NOGs) in *Caulobacterales* species genome representatives of the "core dataset" (emapper v2.1.5). For each genome, attributes are separated by "@" as presented in **Supplementary Data S4**.

**Data S8 | Overview of eggNOG-mapper-predicted KOs.** Overview of eggNOG-mapper[32] (emapper) KEGG ortholog (KO) annotations of **(a)** *Caulobacterales* species genome representatives of the "core dataset" (emapper v2.1.5), **(b)** *Acaudatibacter* gen. nov. ("Palsa-881") genomes of the "extended dataset" (emapper v2.1.12), or **(c)** genomes of curved crescentin-encoding *Hyphomicrobiales* species (emapper v2.1.12). For each genome, attributes are separated by "@" as presented in **Supplementary Data S4**.

**Data S9 | Overview of GhostKOALA-predicted KOs.** Overview of GhostKOALA[38] v2.2 KEGG ortholog (KO) annotations of **(a)** *Caulobacterales* species genome representatives of the "core dataset" (GhostKOALA v2.3), or **(b)** *Acaudatibacter* gen. nov. ("Palsa-881") genomes of the "extended dataset" (GhostKOALA v3.0). For each genome, attributes are separated by "@" as presented in **Supplementary Data S4**.

**Data S10 | Overview of RBHs.** Overview of reciprocal best blast hit (RBH) results for the *C. crescentus* CB15 proteome of the GCF_000006905.1 assembly queried against **(a)** *Caulobacterales* species genome representatives of the "core dataset", and **(b)** *Acaudatibacter* gen. nov. ("Palsa-881") genomes of the "extended dataset". For each genome, attributes are separated by "@" as presented in **Supplementary Data S4**.

**Data S11 | Pairwise ANI.** Overview of pairwise average nucleotide identity (ANI) comparisons using FastANI[145] v1.33 for *Caulobacterales* genomes of the "core dataset" as well as *Acaudatibacter* gen. nov. ("Palsa-881") species of the "extended dataset". For each genome, attributes are separated by "@" as presented in **Supplementary Data S4**. Note that FastANI simply outputs "NA" for ANIs far below 80%.

**Data S12 | Pairwise AAI.** Overview of pairwise average amino acid identity (AAI) comparisons using EzAAI[146] v1.2.3 for *Caulobacterales* genomes of the "core dataset" as well as *Acaudatibacter* gen. nov. ("Palsa-881") species of the "extended dataset". For each genome, attributes are separated by "@" as presented in **Supplementary Data S4**.

**Data S13 | KEGG Decoder: pathway completeness.** Pathway completeness prediction using KEGG Decoder[147] v1.3 with GhostKOALA-predicted KOs listed in **Supplementary Data S9**.

**Data S14 | Manual refinement of species phylogeny.** Sequences manually removed when making the manually refined ML species phylogeny presented in **Fig. 1a** and **Supplementary Fig. S1**, which included the removal of putative paralogs, contamination, long-branching, horizontal transfers, and duplicate sequences.
