## Supplementary Figures, excl. Legends for "Widespread potential for phototrophy and convergent reduction of lifecycle complexity in the dimorphic order *Caulobacterales*"

**Fig. S1**

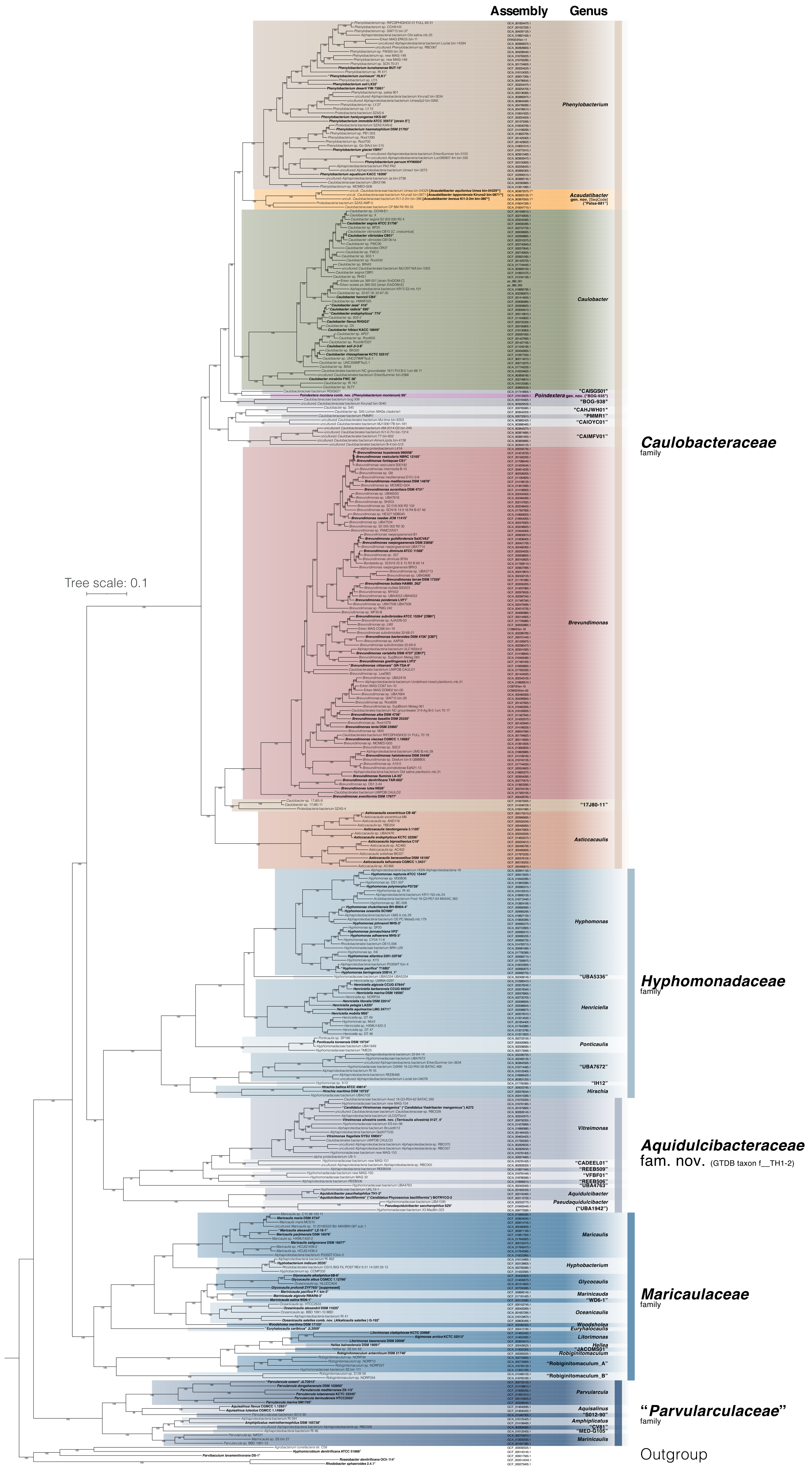
Assembly
Genus
Tree scale: 0.1
Phenylobacterium
Acaudatibacter gen. nov. [SeqCode] ("Palsa-881")
Caulobacter
"CAISGS01"
Poindextera gen. nov. ("BOG-935")
"BOG-938"
"CAHJWH01"
"PMMR1"
"CAIOYC01"
"CAIMFV01"
Caulobacteraceae
family
Brevundimonas
"17J80-11"
Asticcacaulis
Hyphomonas
"UBA5336"
Hyphomonadaceae
family
Henriciella
Ponticaulis
"UBA7672"
"IH12"
Hirschia
Vitreimonas
Aquidulcibacteraceae
fam. nov. (GTDB taxon f__TH1-2)
"CADEEL01"
"REEB509"
"VFBF01"
"REEB506"
"UBA4763"
Aquidulcibacter
Pseudaquidulcibacter
("UBA1942")
Maricaulis
Hyphobacterium
Glycocaulis
Marinicauda
"WD6-1"
Oceanicaulis
Maricaulaceae
family
Woodsholea
Euryhalocaulis
Litorimonas
Hellea
"JACOMS01"
Robiginitomaculum
"Robiginitomaculum_A"
"Robiginitomaculum_B"
Parvularcula
"Parvularculaceae"
family
Aquisalinus
"S012-90"
Amphiplicatus
"CV81"
"MED-G105"
Marinicauda
Outgroup

Fig. S2

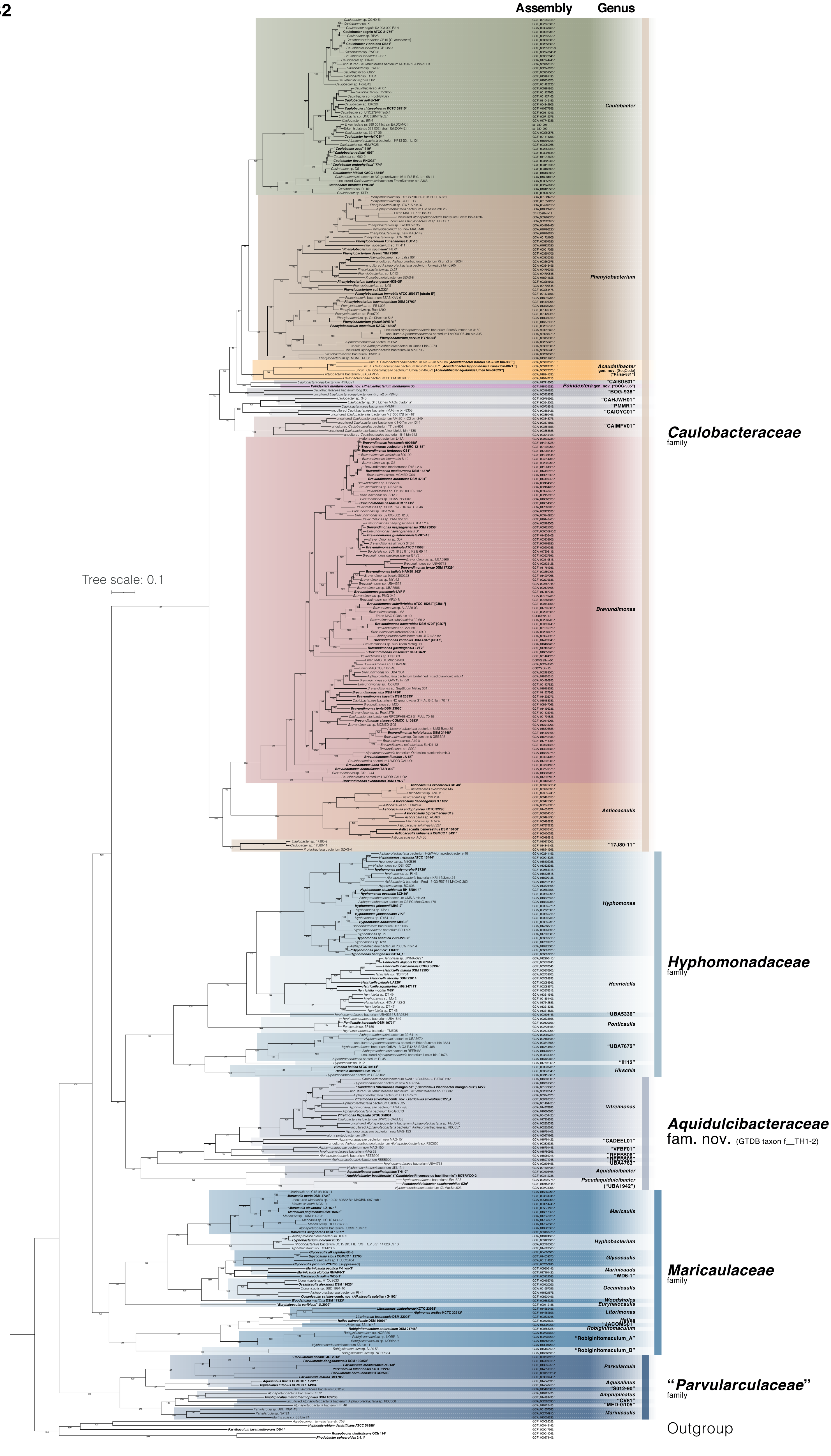

Assembly
Genus
Caulobacter
Phenylobacterium
Acaudatibacter gen. nov. [SeqCode] ("Palsa-881")
Poindextera gen. nov. ("BOG-935")
"CAISGS01"
"BOG-938"
"CAHJWH01"
"PMMR1"
"CAIOYC01"
"CAIMFV01"
Caulobacteraceae
family
Brevundimonas
Asticcacaulis
"17J80-11"
Tree scale: 0.1
Hyphomonas
Hyphomonadaceae
family
Henriciella
"UBA5336"
Ponticaulis
"UBA7672"
"IH12"
Hirschia
Vitreimonas
Aquidulcibacteraceae
fam. nov. (GTDB taxon f__TH1-2)
"CADEEL01"
"VFBF01"
"REEB506"
"REEB509"
"UBA4763"
Aquidulcibacter
Pseudaquidulcibacter ("UBA1942")
Maricaulis
Hyphobacterium
Glycocaulis
Marinicauda
"WD6-1"
Maricaulaceae
family
Oceanicaulis
Woodsholea
Euryhalocaulis
Litorimonas
Hellea
"JACOMS01"
Robiginitomaculum
"Robiginitomaculum_A"
"Robiginitomaculum_B"
Parvularcula
"Parvularculaceae"
family
Aquisalinus
"S012-90"
Amphiplicatus
"CV81"
"MED-G105"
Marinicaulis
Outgroup

**Fig. S3**

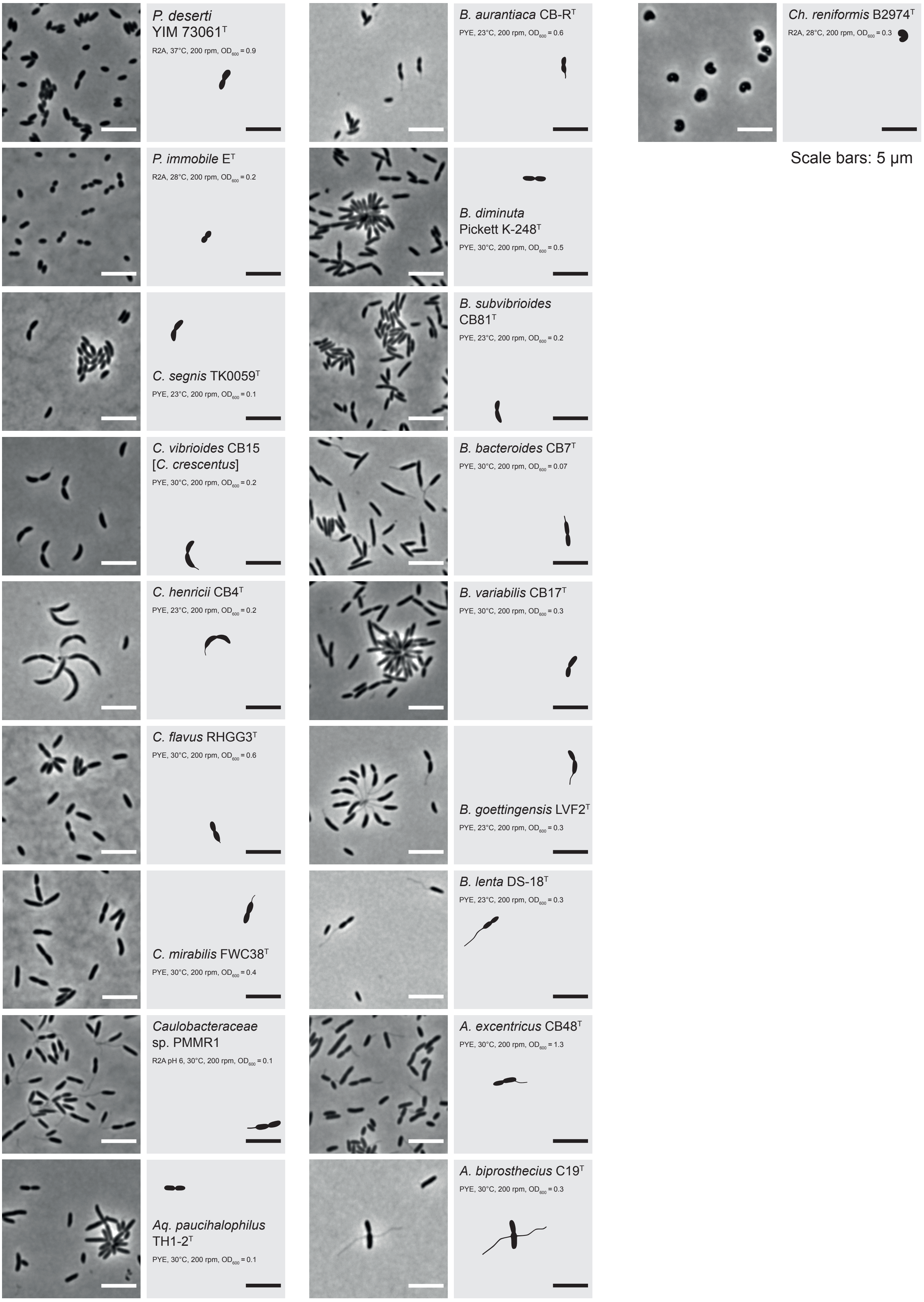

*P. deserti* YIM 73061[T]
R2A, 37°C, 200 rpm, $OD_{600}$ = 0.9
*P. immobile* E[T]
R2A, 28°C, 200 rpm, $OD_{600}$ = 0.2
*C. segnis* TK0059[T]
PYE, 23°C, 200 rpm, $OD_{600}$ = 0.1
*C. vibrioides* CB15 [*C. crescentus*]
PYE, 30°C, 200 rpm, $OD_{600}$ = 0.2
*C. henricii* CB4[T]
PYE, 23°C, 200 rpm, $OD_{600}$ = 0.2
*C. flavus* RHGG3[T]
PYE, 30°C, 200 rpm, $OD_{600}$ = 0.6
*C. mirabilis* FWC38[T]
PYE, 30°C, 200 rpm, $OD_{600}$ = 0.4
*Caulobacteraceae* sp. PMMR1
R2A pH 6, 30°C, 200 rpm, $OD_{600}$ = 0.1
*Aq. paucihalophilus* TH1-2[T]
PYE, 30°C, 200 rpm, $OD_{600}$ = 0.1
*B. aurantiaca* CB-R[T]
PYE, 23°C, 200 rpm, $OD_{600}$ = 0.6
*B. diminuta* Pickett K-248[T]
PYE, 30°C, 200 rpm, $OD_{600}$ = 0.5
*B. subvibrioides* CB81[T]
PYE, 23°C, 200 rpm, $OD_{600}$ = 0.2
*B. bacteroides* CB7[T]
PYE, 30°C, 200 rpm, $OD_{600}$ = 0.07
*B. variabilis* CB17[T]
PYE, 30°C, 200 rpm, $OD_{600}$ = 0.3
*B. goettingensis* LVF2[T]
PYE, 23°C, 200 rpm, $OD_{600}$ = 0.3
*B. lenta* DS-18[T]
PYE, 23°C, 200 rpm, $OD_{600}$ = 0.3
*A. excentricus* CB48[T]
PYE, 30°C, 200 rpm, $OD_{600}$ = 1.3
*A. biprosthecus* C19[T]
PYE, 30°C, 200 rpm, $OD_{600}$ = 0.3
*Ch. reniformis* B2974[T]
R2A, 28°C, 200 rpm, $OD_{600}$ = 0.3
Scale bars: 5 µm

**Fig. S4**

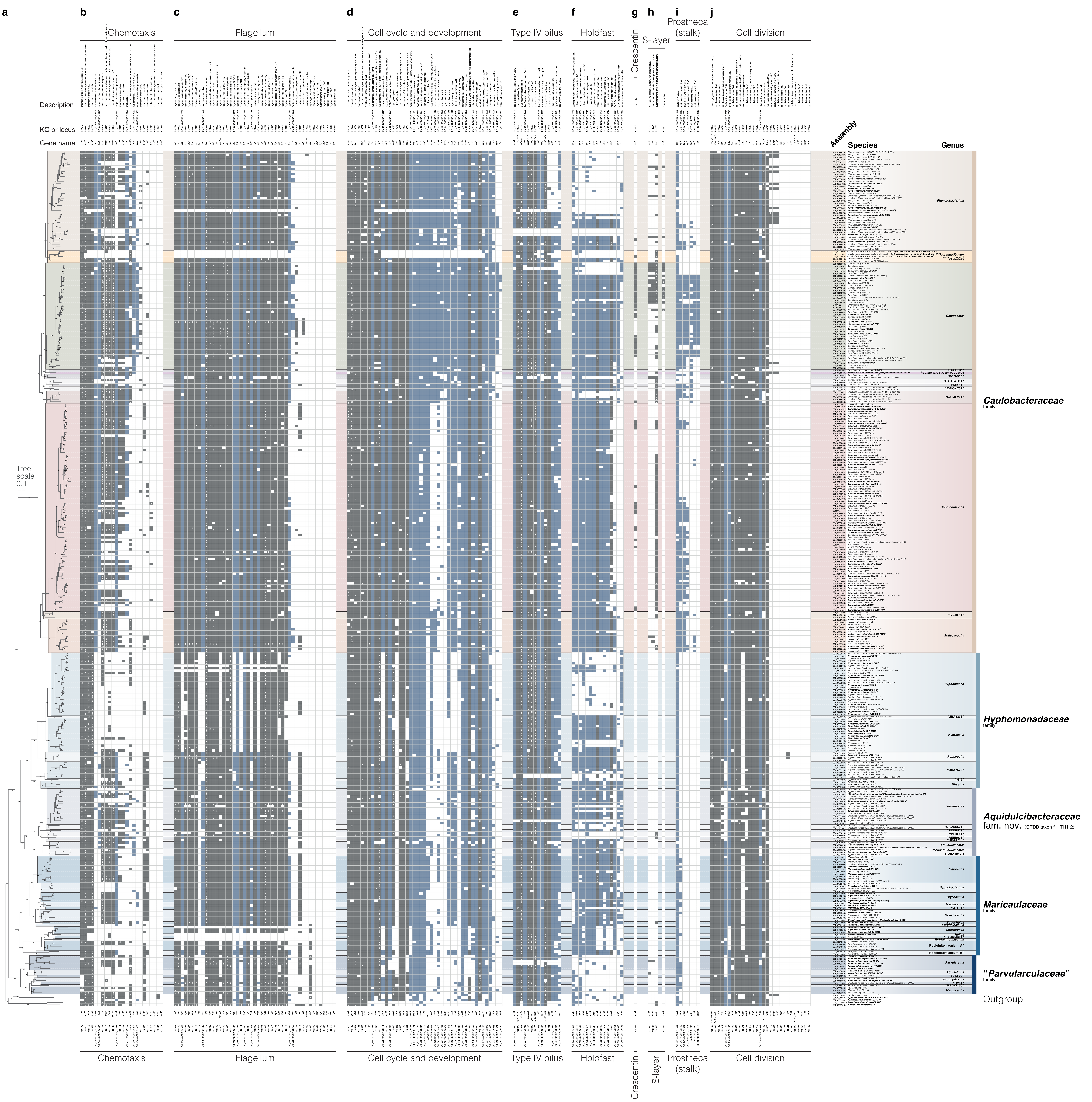
a
b
Chemotaxis
c
Flagellum
d
Cell cycle and development
e
Type IV pilus
f
Holdfast
g
Crescentin
h
S-layer
i
Prostheca (stalk)
j
Cell division
Description
KO or locus
Gene name
Assembly
Species
Genus
Tree scale 0.1
Phenylobacterium
Caulobacter
Brevundimonas
Asticcacaulis
Hyphomonas
Henriciella
Ponticaulis
Hirschia
Vitreimonas
Maricaulis
Hyphobacterium
Glycocaulis
Oceanicaulis
Parvularcula
Caulobacteraceae
family
Hyphomonadaceae
family
Aquidulcibacteraceae
fam. nov. (GTDB taxon f__TH1-2)
Maricaulaceae
family
“Parvularculaceae”
family
Outgroup

### Fig. S5

**a**

#### *Acaudatibacter* (GTDB taxon g__Palsa-881) species clusters

| *Acaudatibacter* (GTDB taxon g__Palsa-881) species clusters | completeness (%) | contamination (%) | assembly size (bp) | estimated genome size (bp) | G+C content | no. contigs |
|---|---|---|---|---|---|---|
| ***Ac. boreus*** **(sp. 21122) [n = 4]:** | | | | | | |
| **uncultured *Caulobacteraceae* bacterium Ki1-1-6m_bin-0605**: GCA_903839265.1 | 94.32 | 3.83 | 4 342 596 | 4 604 311 | 67.51% | 610 |
| * **uncultured *Caulobacteraceae* bacterium Ki1-2-2m_bin-386$^{Ts}$**: GCA_903870555.1$^{Ts}$ | 94.46 | 2.03 | 4 349 168 | 4 604 298 | 67.46% | 692 |
| Ki1-1-6m_bin-20 | 93.45 | 3.32 | 4 228 465 | 4 524 839 | 67.60% | 576 |
| Ki1-2-2m_bin-14 | 94.46 | 2.03 | 4 362 619 | 4 618 538 | 67.51% | 699 |
| ***Ac. lapponiensis*** **(sp. 21175) [n = 4]:** | | | | | | |
| **uncultured *Caulobacteraceae* bacterium Ki2-D3_bin-0046**: GCA_903840985.1 | 96.97 | 2.32 | 4 624 202 | 4 768 639 | 68.83% | 304 |
| Ki2-D3_bin-69 | 96.97 | 1.94 | 4 584 716 | 4 727 920 | 68.84% | 288 |
| **uncultured *Caulobacteraceae* bacterium Kiruna_bin-06165**: GCA_903860905.1 | 94.86 | 4.47 | 4 566 348 | 4 813 768 | 68.84% | 357 |
| * **uncultured *Caulobacteraceae* bacterium Kiruna2_bin-0871$^{Ts}$**: GCA_903923135.1$^{Ts}$ | 96.47 | 2.20 | 4 636 946 | 4 806 723 | 68.83% | 279 |
| ***Ac. aquilonius*** **(sp.19747) [n = 12]:** | | | | | | |
| **uncultured *Caulobacteraceae* bacterium AlinenLipids_bin-4060**: GCA_903916695.1 | 94.29 | 2.54 | 4 322 818 | 4 584 462 | 67.43% | 238 |
| AM-lipid-02-D3_bin-31 | 69.50 | 2.22 | 3 060 937 | 4 404 258 | 67.58% | 950 |
| AM-lipid-02-D3_megahit_metabat_bin-0449 | 69.71 | 2.68 | 3 125 578 | 4 483 652 | 67.57% | 973 |
| **uncultured *Caulobacteraceae* bacterium KR_bin-3969**: GCA_903911855.1 | 94.84 | 3.14 | 4 415 891 | 4 656 142 | 67.38% | 476 |
| **uncultured *Caulobacteraceae* bacterium KR2_bin-0350**: GCA_903841115.1 | 91.50 | 4.21 | 4 149 148 | 4 534 617 | 67.41% | 675 |
| KR2_bin-4 | 91.15 | 3.23 | 4 027 975 | 4 418 912 | 67.44% | 637 |
| * **uncultured *Caulobacteraceae* bacterium Umea_bin-04329$^{Ts}$**: GCA_903872075.1$^{Ts}$ | 95.32 | 1.62 | 4 361 582 | 4 575 761 | 67.43% | 100 |
| **uncultured *Caulobacteraceae* bacterium Umea3p1_bin-1515**: GCA_903876365.1 | 97.53 | 3.43 | 4 556 442 | 4 671 627 | 67.37% | 206 |
| **uncultured *Caulobacteraceae* bacterium Umea3_bin-1590**: GCA_903887165.1 | 95.59 | 2.45 | 4 477 020 | 4 683 615 | 67.42% | 226 |
| Umea3p1_bin-46 | 97.53 | 1.93 | 4 390 252 | 4 501 236 | 67.43% | 164 |
| **uncultured *Caulobacteraceae* bacterium VK_bin-1681**: GCA_903891275.1 | 94.67 | 3.05 | 4 268 743 | 4 509 001 | 67.42% | 502 |
| VK3_bin-17 | 86.78 | 2.38 | 3 837 298 | 4 421 918 | 67.48% | 810 |
| ***Ac.* sp. bog_900** (sp. 18536) [n = 2]: | | | | | | |
| ***Alphaproteobacteria* bog_909**: GCA_003133765.1 | 82.74 | 1.98 | 2 735 410 | 3 305 832 | 65.34% | 333 |
| ***Alphaproteobacteria* bog_900**: GCA_003136615.1 | 86.13 | 2.15 | 2 852 624 | 3 312 124 | 65.37% | 311 |
| ***Ac.* sp. CP_BE_RX_R2_21** (sp. 19682) [n = 1]: ***Caulobacteraceae* bacterium CP_BE_RX_R2_21**: GCA_019232665.1 | 93.53 | 2.65 | 3 828 023 | 4 092 674 | 67.38% | 375 |
| ***Ac.* sp. 3300015024_21** (sp. 06679) [n = 1]: 3300015024_21 | 96.08 | 3.86 | 3 843 261 | 3 999 970 | 64.12% | 241 |
| ***Ac.* sp. palsa_881** (sp. 18537) [n = 1]: ***Alphaproteobacteria* palsa_881**: GCA_003161535.1 | 71.05 | 2.52 | 1 908 427 | 2 685 984 | 67.25% | 414 |
| ***Ac.* sp. 3300025574_7** (sp. 11035) [n = 1]: 3300025574_7 | 97.71 | 1.13 | 3 193 443 | 3 268 433 | 68.05% | 89 |
| ***Ac.* sp. MGR_bin405** (sp. 19143) [n = 1]: ***Caulobacteraceae* bacterium MGR_bin405**: GCA_013815405.1 | 79.31 | 2.61 | 2 720 259 | 3 429 787 | 68.56% | 527 |
| ***Ac.* sp. Umea1p3_bin-3** (sp. 23796) [n = 5]: | | | | | | |
| Kiruna_megahit_metabat_bin-10665 | 67.32 | 4.17 | 3 041 251 | 4 517 324 | 63.10% | 885 |
| Umea1p2_bin-48 | 50.98 | 0.97 | 2 061 914 | 4 044 360 | 63.50% | 798 |
| Umea1p2_megahit_metabat_bin-0087 | 53.32 | 4.22 | 2 411 118 | 4 521 853 | 63.40% | 935 |
| Umea1p3_bin-3 | 92.53 | 3.33 | 4 063 138 | 4 391 040 | 63.19% | 642 |
| VK1_bin-18 | 80.40 | 1.10 | 3 420 437 | 4 254 018 | 63.34% | 904 |
| * ***Ac.* sp. SZAS AMP-5** (sp. 19591) [n = 1]: ***Proteobacteria* bacterium SZAS_AMP-5**: GCA_018241335.1 | 98.27 | 2.46 | 4 405 988 | 4 483 627 | 70.14% | 458 |
| * ***Ac.* sp. CP_BM_RX_R9_33** (sp. 19683) [n = 1]: ***Caulobacteraceae* bacterium CP_BM_RX_R9_33**: GCA_019247715.1 | 97.86 | 3.13 | 4 056 632 | 4 145 317 | 69.93% | 108 |

**b**

##### Sampling environment metadata

| Geographic location | Lv1 | Lv2 | Lv3 | Depth (m) | $O_2$ (mg/L) | Temp. (°C) | pH |
|---|---|---|---|---|---|---|---|
| **Sweden**: Kiruna, Lake Ki1 | | | lake water | 1.6 | 0.3 | 16.1 | 4.99 |
| **Sweden**: Kiruna, Lake Ki1 | | | lake water | 2.2 | -0.1 | 9.5 | 5.19 |
| **Sweden**: Kiruna, Lake Ki1 | | | lake water | 1.6 | 0.3 | 16.1 | 4.99 |
| **Sweden**: Kiruna, Lake Ki1 | | | lake water | 2.2 | -0.1 | 9.5 | 5.19 |
| **Sweden**: Kiruna, Lake Ki2 | | | lake water | 4.2 | 0 | 7.4 | 5.01 |
| **Sweden**: Kiruna, Lake Ki2 | | | lake water | 4.2 | 0 | 7.4 | 5.01 |
| **Sweden**: Kiruna, Lake Ki2 | | | lake water | | | | |
| **Sweden**: Kiruna, Lake Ki2 | | | lake water | | | | |
| **Finland**: Evo, Alinen Mustajärvi | | | lake water | | | | |
| **Finland**: Evo, Alinen Mustajärvi | | | lake water | 2.3 | 1.55 | 12.5 | |
| **Finland**: Evo, Alinen Mustajärvi | | | lake water | 2.3 | 1.55 | 12.5 | |
| **Finland**: Evo, Keskinen Rajajärvi | | | lake water | | | | |
| **Finland**: Evo, Keskinen Rajajärvi | | | lake water | 2 | 1.5 | 12.2 | |
| **Finland**: Evo, Keskinen Rajajärvi | | | lake water | 2 | 1.5 | 12.2 | |
| **Sweden**: Umeå | | | lake water | | | | |
| **Sweden**: Umeå, Björntjärnen | | | lake water | 0.25 | 8.7 | 13.8 | 5.38 |
| **Sweden**: Umeå, Björntjärnen | | | lake water | | | | |
| **Sweden**: Umeå, Björntjärnen | | | lake water | 0.25 | 8.7 | 13.8 | 5.38 |
| **Finland**: Evo, Valkea Kotinen | | | lake water | | | | |
| **Finland**: Evo, Valkea Kotinen | | | lake water | 3.25 | 0.46 | 9.8 | |
| **Sweden**: Stordalen Mire | | | permafrost active layer soil | | | | |
| **Sweden**: Stordalen Mire | | | permafrost active layer soil | | | | |
| **Brazil**: Minas Gerais | | | rhizosphere of *Vellozia epidendroides* | | | | |
| **Sweden**: Tarfala | | | glacier forefield soil | | | | |
| **Sweden**: Stordalen Mire | | | permafrost active layer soil | | | | |
| **USA**: Alaska, Barrow | | | permafrost | | | | |
| **Antarctica**: Mackay Glacier reg. | | | cold desert soils from glacier region | | | | |
| **Sweden**: Kiruna | | | lake water | | | | |
| **Sweden**: Umeå, Nästjärnen | | | lake water | 3.5 | 0.5 | 10.9 | 5.67 |
| **Sweden**: Umeå, Nästjärnen | | | lake water | 3.5 | 0.5 | 10.9 | 5.67 |
| **Sweden**: Umeå, Nästjärnen | | | lake water | 4 | -0.1 | 8.5 | 5.46 |
| **Finland**: Evo, Valkea Kotinen | | | lake water | 0.5 | 8.47 | 19.5 | |
| **China**: Shenzhen | | | activated sludge | | | | |
| **Brazil**: Minas Gerais | | | rhizosphere of *Barbacenia macrantha* | | | | |

**c**

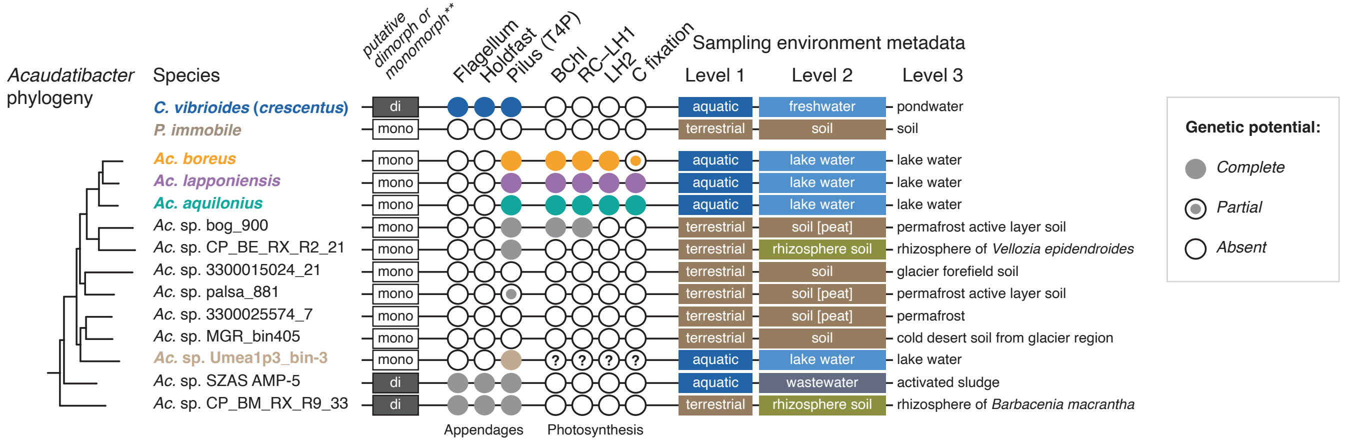

### Fig. S6

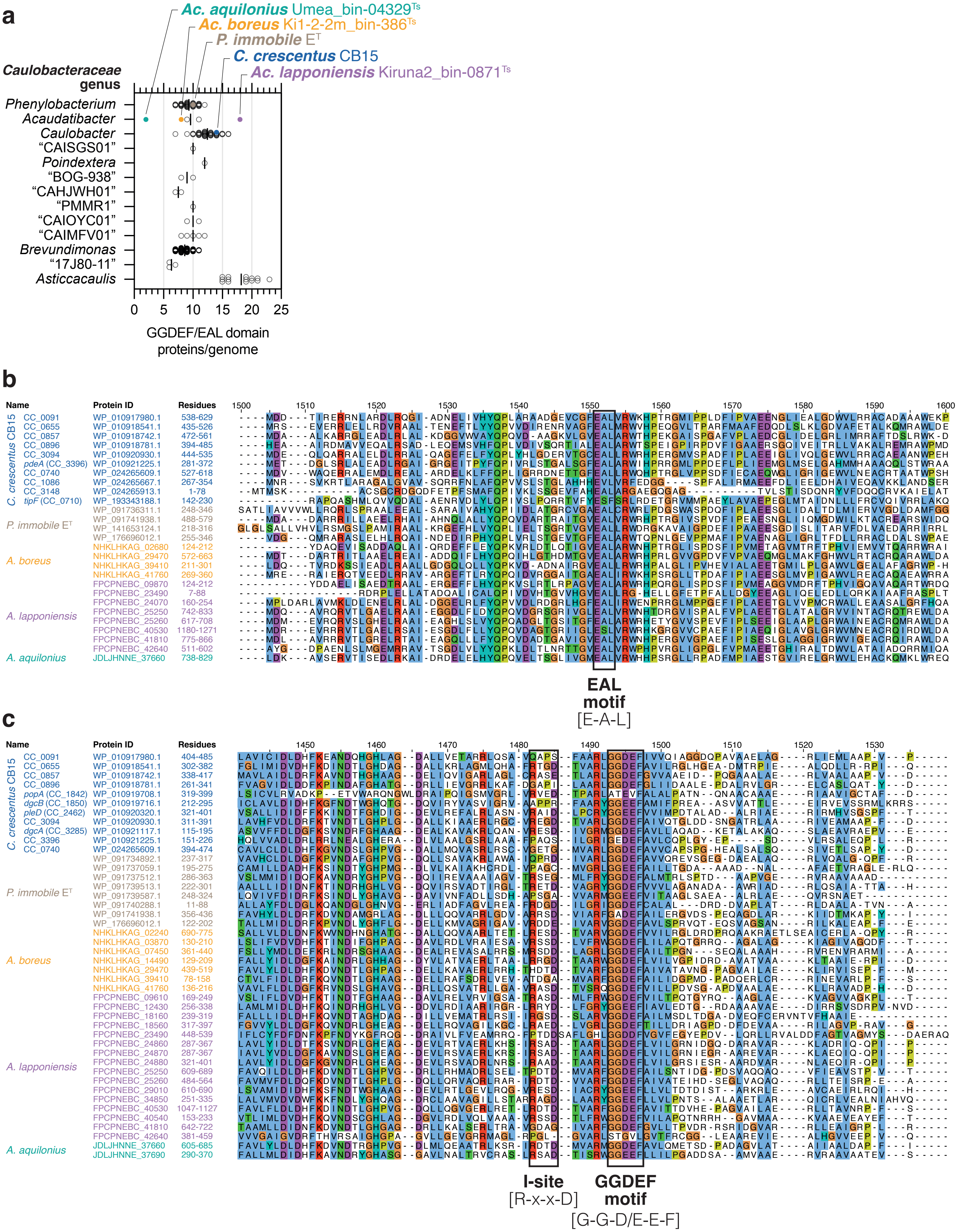
a
Ac. aquilonius Umea_bin-04329Ts
Ac. boreus Ki1-2-2m_bin-386Ts
P. immobile ET
C. crescentus CB15
Ac. lapponiensis Kiruna2_bin-0871Ts
Caulobacteraceae genus
Phenylobacterium
Acaudatibacter
Caulobacter
"CAISGS01"
Poindextera
"BOG-938"
"CAHJWH01"
"PMMR1"
"CAIOYC01"
"CAIMFV01"
Brevundimonas
"17J80-11"
Asticcacaulis
0
5
10
15
20
25
GGDEF/EAL domain proteins/genome
b
Name
Protein ID
Residues
C. crescentus CB15
P. immobile ET
A. boreus
A. lapponiensis
A. aquilonius
EAL motif
[E-A-L]
c
Name
Protein ID
Residues
I-site
[R-x-x-D]
GGDEF motif
[G-G-D/E-E-F]

**Fig. S7**

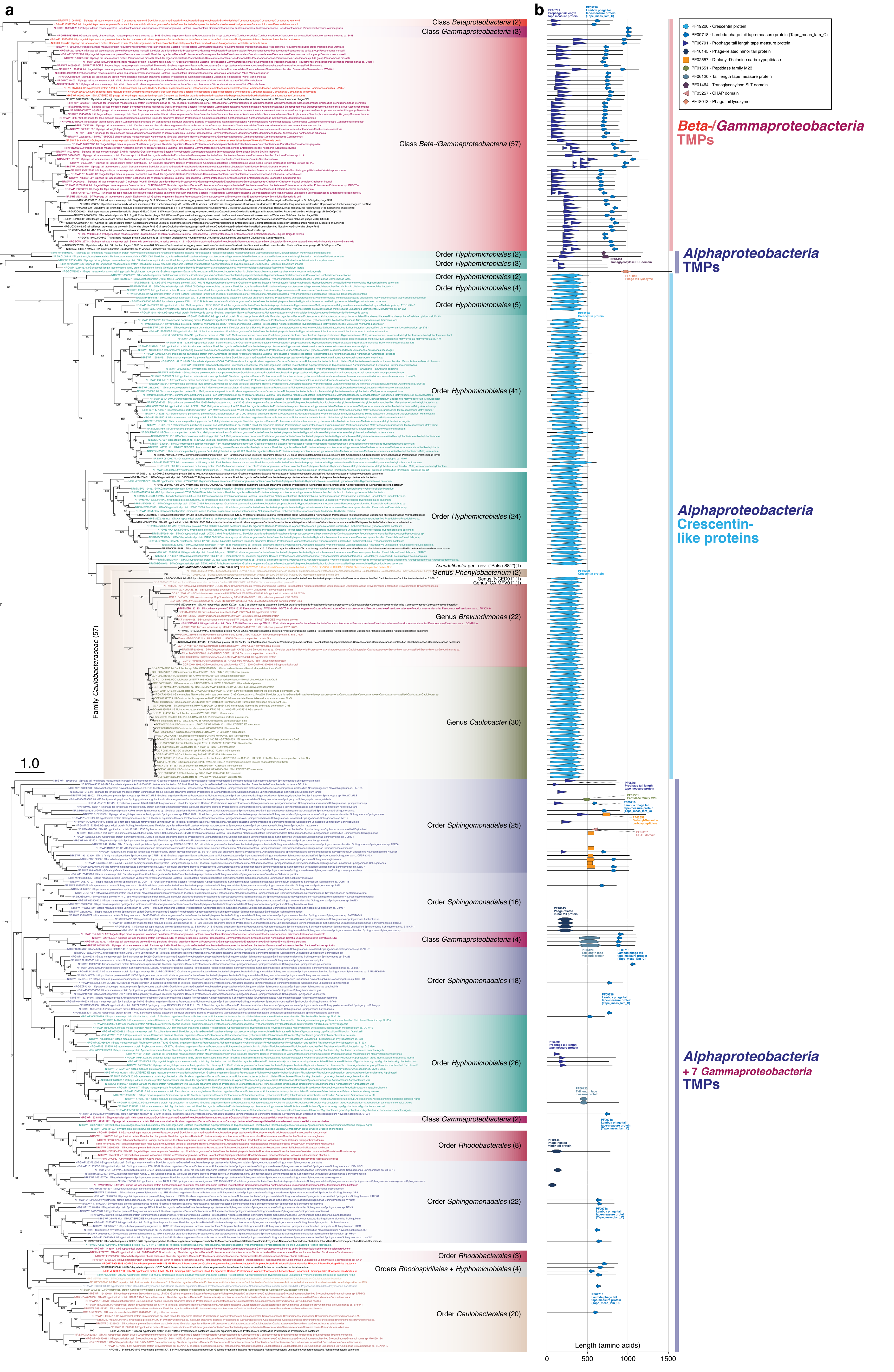
a
b
Class Betaproteobacteria (2)
Class Gammaproteobacteria (3)
Class Beta-/Gammaproteobacteria (57)
Order Hyphomicrobiales (2)
Order Hyphomicrobiales (3)
Order Hyphomicrobiales (2)
Order Hyphomicrobiales (4)
Order Hyphomicrobiales (5)
Order Hyphomicrobiales (41)
Order Hyphomicrobiales (24)
Acaudatibacter gen. nov. ("Palsa-881")(1)
Genus Phenylobacterium (2)
Genus "NCED01" (1)
Genus "CAIMFV01" (1)
Genus Brevundimonas (22)
Genus Caulobacter (30)
Family Caulobacteraceae (57)
1.0
Order Sphingomonadales (25)
Order Sphingomonadales (16)
Class Gammaproteobacteria (4)
Order Sphingomonadales (18)
Order Hyphomicrobiales (26)
Class Gammaproteobacteria (2)
Order Rhodobacterales (8)
Order Sphingomonadales (22)
Order Rhodobacterales (3)
Orders Rhodospirillales + Hyphomicrobiales (4)
Order Caulobacterales (20)
PF19220 - Crescentin protein
PF09718 - Lambda phage tail tape-measure protein (Tape_meas_lam_C)
PF06791 - Prophage tail length tape measure protein
PF10145 - Phage-related minor tail protein
PF02557 - D-alanyl-D-alanine carboxypeptidase
PF01551 - Peptidase family M23
PF06120 - Tail length tape measure protein
PF01464 - Transglycosylase SLT domain
PF05257 - CHAP domain
PF18013 - Phage tail lysozyme
Beta-/Gammaproteobacteria TMPs
Alphaproteobacteria TMPs
Alphaproteobacteria Crescentin-like proteins
Alphaproteobacteria + 7 Gammaproteobacteria TMPs
Length (amino acids)
0
500
1000
1500

#### Fig. S8

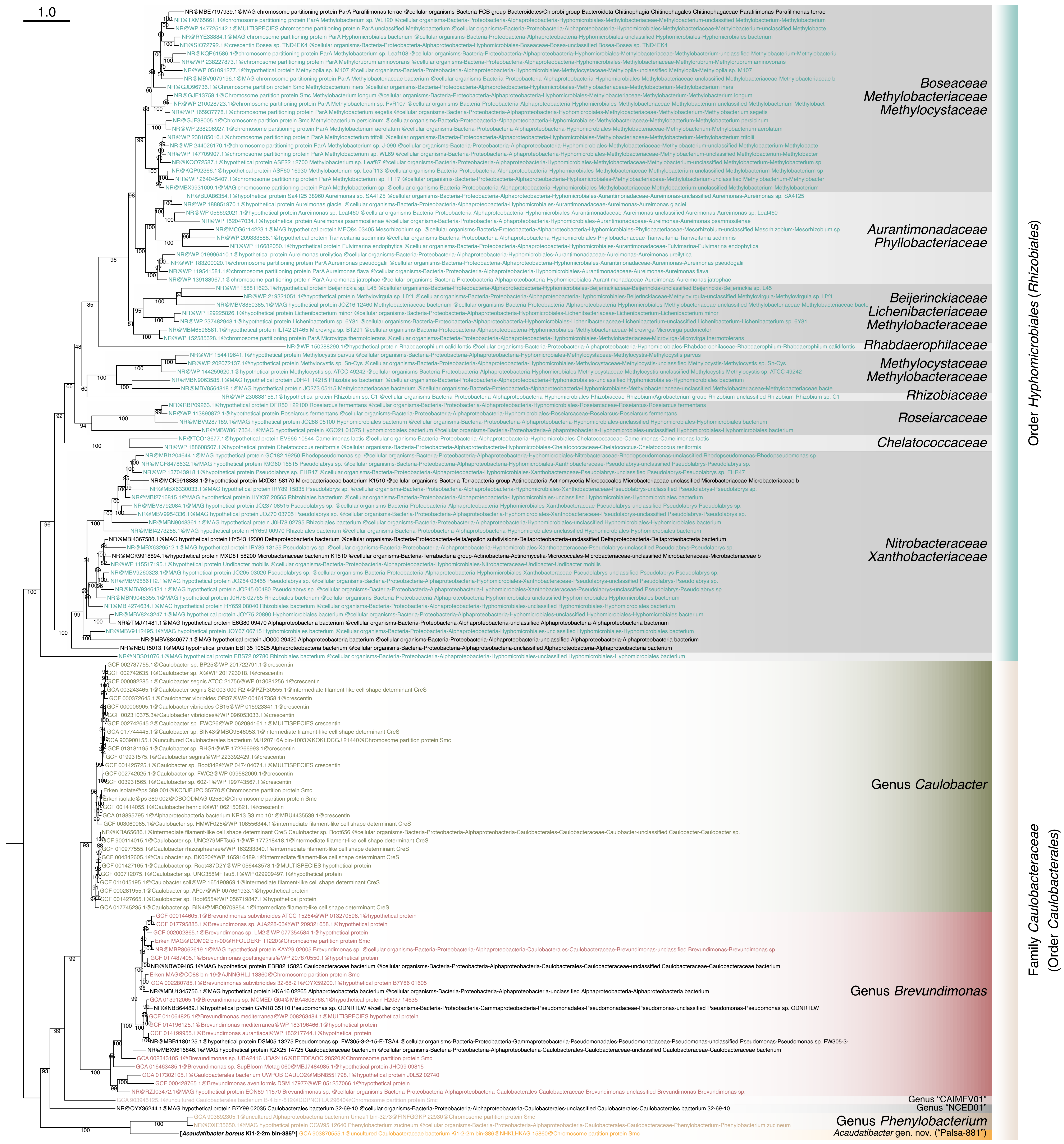
1.0
Boseaceae
Methylobacteriaceae
Methylocystaceae
Aurantimonadaceae
Phyllobacteriaceae
Beijerinckiaceae
Lichenibacteriaceae
Methylobacteriaceae
Rhabdaerophilaceae
Methylocystaceae
Methylobacteriaceae
Rhizobiaceae
Roseiarcaceae
Chelatococcaceae
Nitrobacteraceae
Xanthobacteriaceae
Order Hyphomicrobiales (Rhizobiales)
Genus Caulobacter
Genus Brevundimonas
Genus "CAIMFV01"
Genus "NCED01"
Genus Phenylobacterium
Acaudatibacter gen. nov. ("Palsa-881")
Family Caulobacteraceae
(Order Caulobacterales)

#### Fig. S9

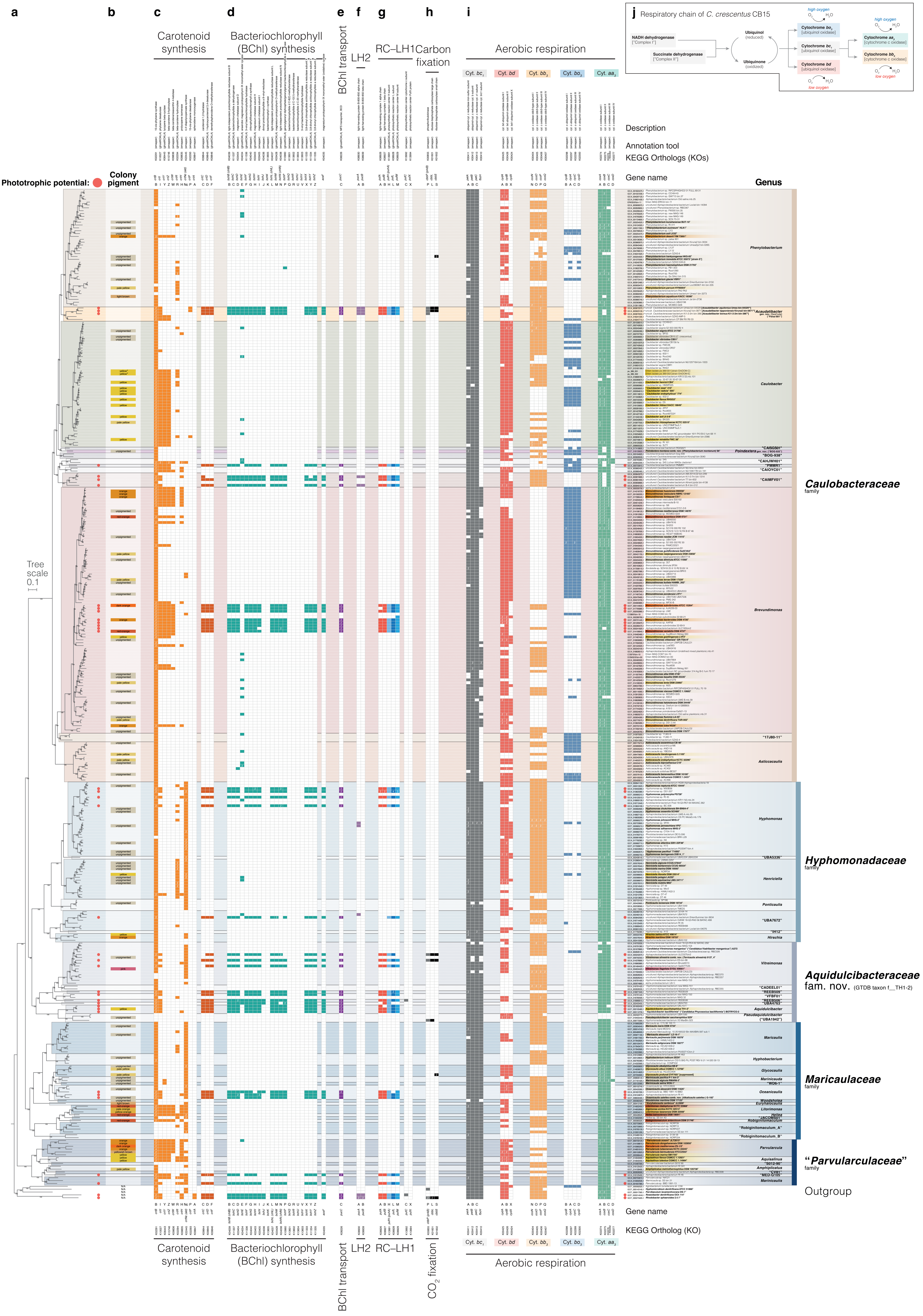

a
b
c
d
e
f
g
h
i
j
Carotenoid synthesis
Bacteriochlorophyll (BChl) synthesis
BChl transport
LH2
RC–LH1
Carbon fixation
Aerobic respiration
Cyt. bc1
Cyt. bd
Cyt. bb3
Cyt. bo3
Cyt. aa3
Respiratory chain of C. crescentus CB15
NADH dehydrogenase ["Complex I"]
Succinate dehydrogenase ["Complex II"]
Ubiquinol (reduced)
Ubiquinone (oxidized)
Cytochrome bo3 [ubiquinol oxidase]
Cytochrome bc1 [ubiquinol oxidase]
Cytochrome bd [ubiquinol oxidase]
Cytochrome aa3 [cytochrome c oxidase]
Cytochrome bb3 [cytochrome c oxidase]
high oxygen
low oxygen
Description
Annotation tool
KEGG Orthologs (KOs)
Gene name
Genus
Phototrophic potential:
Colony pigment
Tree scale 0.1
Phenylobacterium
Acaudatibacter
Caulobacter
Poindextera
Brevundimonas
Asticcacaulis
Hyphomonas
Henriciella
Ponticaulis
Hirschia
Vitreimonas
Aquidulcibacter
Pseudaquidulcibacter
Maricaulis
Hyphobacterium
Glycocaulis
Marinicauda
Oceanicaulis
Woodsholea
Euryhalocaulis
Litorimonas
Hellea
Robiginitomaculum
Parvularcula
Aquisalinus
Amphiplicatus
Caulobacteraceae family
Hyphomonadaceae family
Aquidulcibacteraceae fam. nov. (GTDB taxon f__TH1-2)
Maricaulaceae family
"Parvularculaceae" family
Outgroup
Gene name
KEGG Ortholog (KO)
CO2 fixation

### Fig. S10

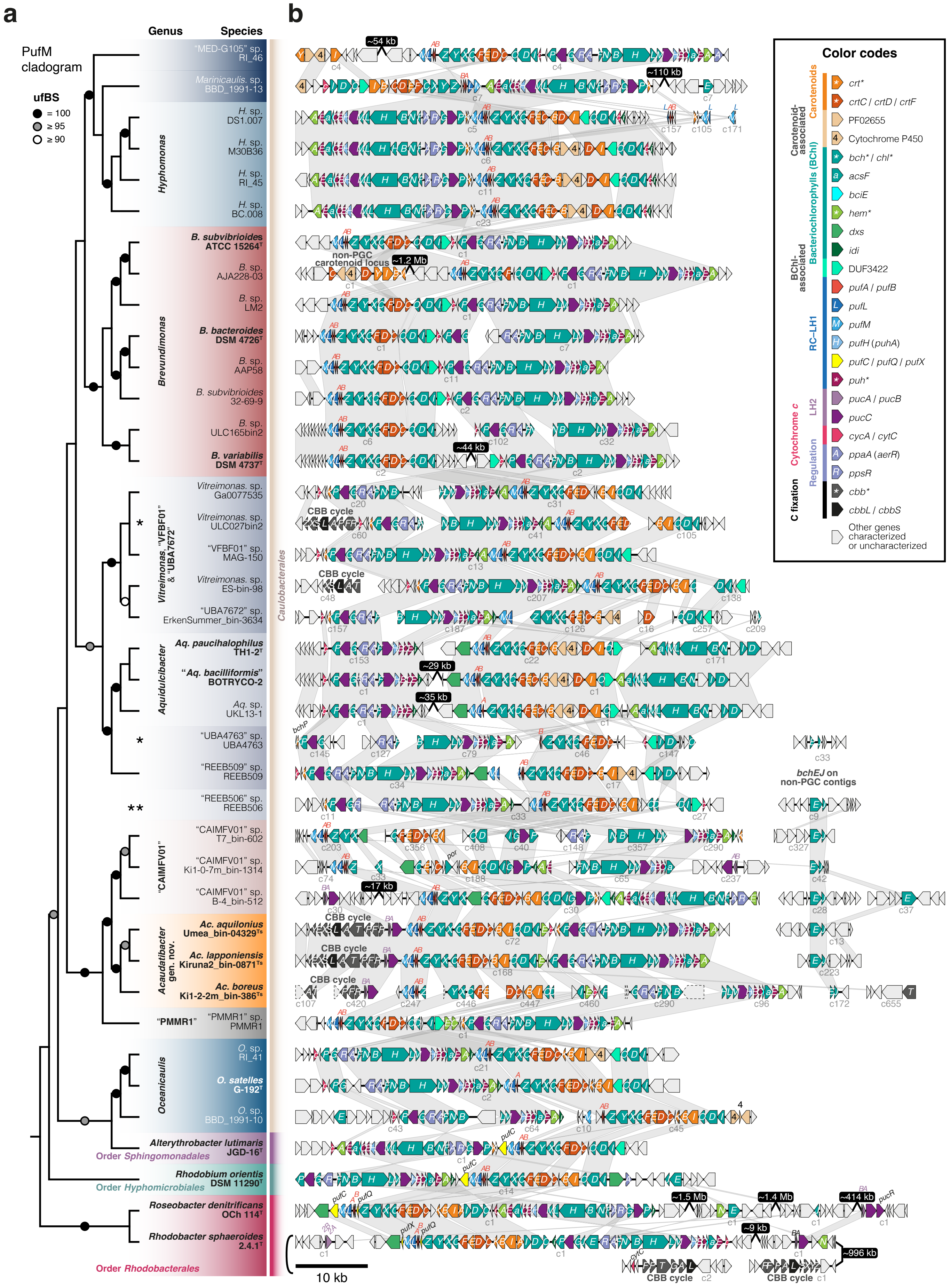
a
b
PufM cladogram
ufBS
= 100
≥ 95
≥ 90
Genus
Species
"MED-G105" sp. RI_46
Marinicaulis. sp. BBD_1991-13
Hyphomonas
H. sp. DS1.007
H. sp. M30B36
H. sp. RI_45
H. sp. BC.008
Brevundimonas
B. subvibrioides ATCC 15264T
B. sp. AJA228-03
B. sp. LM2
B. bacteroides DSM 4726T
B. sp. AAP58
B. subvibrioides 32-69-9
B. sp. ULC165bin2
B. variabilis DSM 4737T
Vitreimonas, "VFBF01" & "UBA7672"
Vitreimonas. sp. Ga0077535
Vitreimonas. sp. ULC027bin2
"VFBF01" sp. MAG-150
Vitreimonas. sp. ES-bin-98
"UBA7672" sp. ErkenSummer_bin-3634
Aquidulcibacter
Aq. paucihalophilus TH1-2T
"Aq. bacilliformis" BOTRYCO-2
Aq. sp. UKL13-1
"UBA4763" sp. UBA4763
"REEB509" sp. REEB509
"REEB506" sp. REEB506
"CAIMFV01"
"CAIMFV01" sp. T7_bin-602
"CAIMFV01" sp. Ki1-0-7m_bin-1314
"CAIMFV01" sp. B-4_bin-512
Acaudatibacter gen. nov.
Ac. aquilonius Umea_bin-04329Ts
Ac. lapponiensis Kiruna2_bin-0871Ts
Ac. boreus Ki1-2-2m_bin-386Ts
"PMMR1"
"PMMR1" sp. PMMR1
Oceanicaulis
O. sp. RI_41
O. satelles G-192T
O. sp. BBD_1991-10
Caulobacterales
Alterythrobacter lutimaris JGD-16T
Order Sphingomonadales
Rhodobium orientis DSM 11290T
Order Hyphomicrobiales
Roseobacter denitrificans OCh 114T
Rhodobacter sphaeroides 2.4.1T
Order Rhodobacterales
~54 kb
~110 kb
~1.2 Mb
non-PGC carotenoid locus
~44 kb
CBB cycle
~29 kb
~35 kb
bchP
bchEJ on non-PGC contigs
~17 kb
~1.5 Mb
~1.4 Mb
~414 kb
~9 kb
~996 kb
10 kb
Color codes
Carotenoids
Carotenoid-associated
crt*
crtC / crtD / crtF
PF02655
Cytochrome P450
Bacteriochlorophylls (BChl)
BChl-associated
bch* / chl*
acsF
bciE
hem*
dxs
idi
DUF3422
RC–LH1
pufA / pufB
pufL
pufM
pufH (puhA)
pufC / pufQ / pufX
puh*
LH2
pucA / pucB
pucC
Cytochrome c
cycA / cytC
Regulation
ppaA (aerR)
ppsR
C fixation
cbb*
cbbL / cbbS
Other genes characterized or uncharacterized

**Fig. S11**

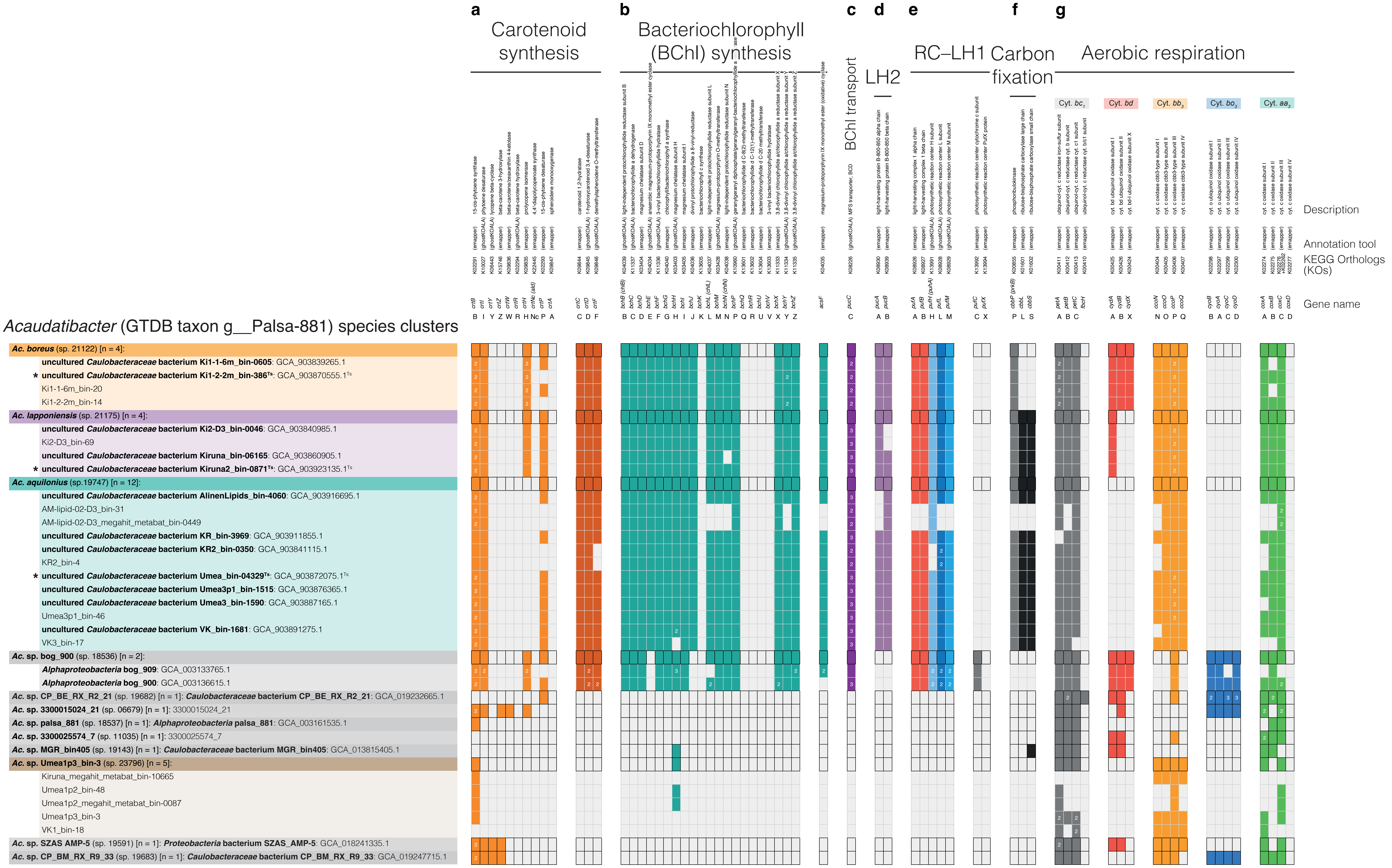
a
Carotenoid synthesis
b
Bacteriochlorophyll (BChl) synthesis
c
BChl transport
d
LH2
e
RC–LH1
f
Carbon fixation
g
Aerobic respiration
Cyt. bc1
Cyt. bd
Cyt. bb3
Cyt. bo3
Cyt. aa3
Description
Annotation tool
KEGG Orthologs (KOs)
Gene name
Acaudatibacter (GTDB taxon g__Palsa-881) species clusters
Ac. boreus (sp. 21122) [n = 4]:
uncultured Caulobacteraceae bacterium Ki1-1-6m_bin-0605: GCA_903839265.1
* uncultured Caulobacteraceae bacterium Ki1-2-2m_bin-386Ts: GCA_903870555.1Ts
Ki1-1-6m_bin-20
Ki1-2-2m_bin-14
Ac. lapponiensis (sp. 21175) [n = 4]:
uncultured Caulobacteraceae bacterium Ki2-D3_bin-0046: GCA_903840985.1
Ki2-D3_bin-69
uncultured Caulobacteraceae bacterium Kiruna_bin-06165: GCA_903860905.1
* uncultured Caulobacteraceae bacterium Kiruna2_bin-0871Ts: GCA_903923135.1Ts
Ac. aquilonius (sp.19747) [n = 12]:
uncultured Caulobacteraceae bacterium AlinenLipids_bin-4060: GCA_903916695.1
AM-lipid-02-D3_bin-31
AM-lipid-02-D3_megahit_metabat_bin-0449
uncultured Caulobacteraceae bacterium KR_bin-3969: GCA_903911855.1
uncultured Caulobacteraceae bacterium KR2_bin-0350: GCA_903841115.1
KR2_bin-4
* uncultured Caulobacteraceae bacterium Umea_bin-04329Ts: GCA_903872075.1Ts
uncultured Caulobacteraceae bacterium Umea3p1_bin-1515: GCA_903876365.1
uncultured Caulobacteraceae bacterium Umea3_bin-1590: GCA_903887165.1
Umea3p1_bin-46
uncultured Caulobacteraceae bacterium VK_bin-1681: GCA_903891275.1
VK3_bin-17
Ac. sp. bog_900 (sp. 18536) [n = 2]:
Alphaproteobacteria bog_909: GCA_003133765.1
Alphaproteobacteria bog_900: GCA_003136615.1
Ac. sp. CP_BE_RX_R2_21 (sp. 19682) [n = 1]: Caulobacteraceae bacterium CP_BE_RX_R2_21: GCA_019232665.1
Ac. sp. 3300015024_21 (sp. 06679) [n = 1]: 3300015024_21
Ac. sp. palsa_881 (sp. 18537) [n = 1]: Alphaproteobacteria palsa_881: GCA_003161535.1
Ac. sp. 3300025574_7 (sp. 11035) [n = 1]: 3300025574_7
Ac. sp. MGR_bin405 (sp. 19143) [n = 1]: Caulobacteraceae bacterium MGR_bin405: GCA_013815405.1
Ac. sp. Umea1p3_bin-3 (sp. 23796) [n = 5]:
Kiruna_megahit_metabat_bin-10665
Umea1p2_bin-48
Umea1p2_megahit_metabat_bin-0087
Umea1p3_bin-3
VK1_bin-18
Ac. sp. SZAS AMP-5 (sp. 19591) [n = 1]: Proteobacteria bacterium SZAS_AMP-5: GCA_018241335.1
Ac. sp. CP_BM_RX_R9_33 (sp. 19683) [n = 1]: Caulobacteraceae bacterium CP_BM_RX_R9_33: GCA_019247715.1

Fig. S12

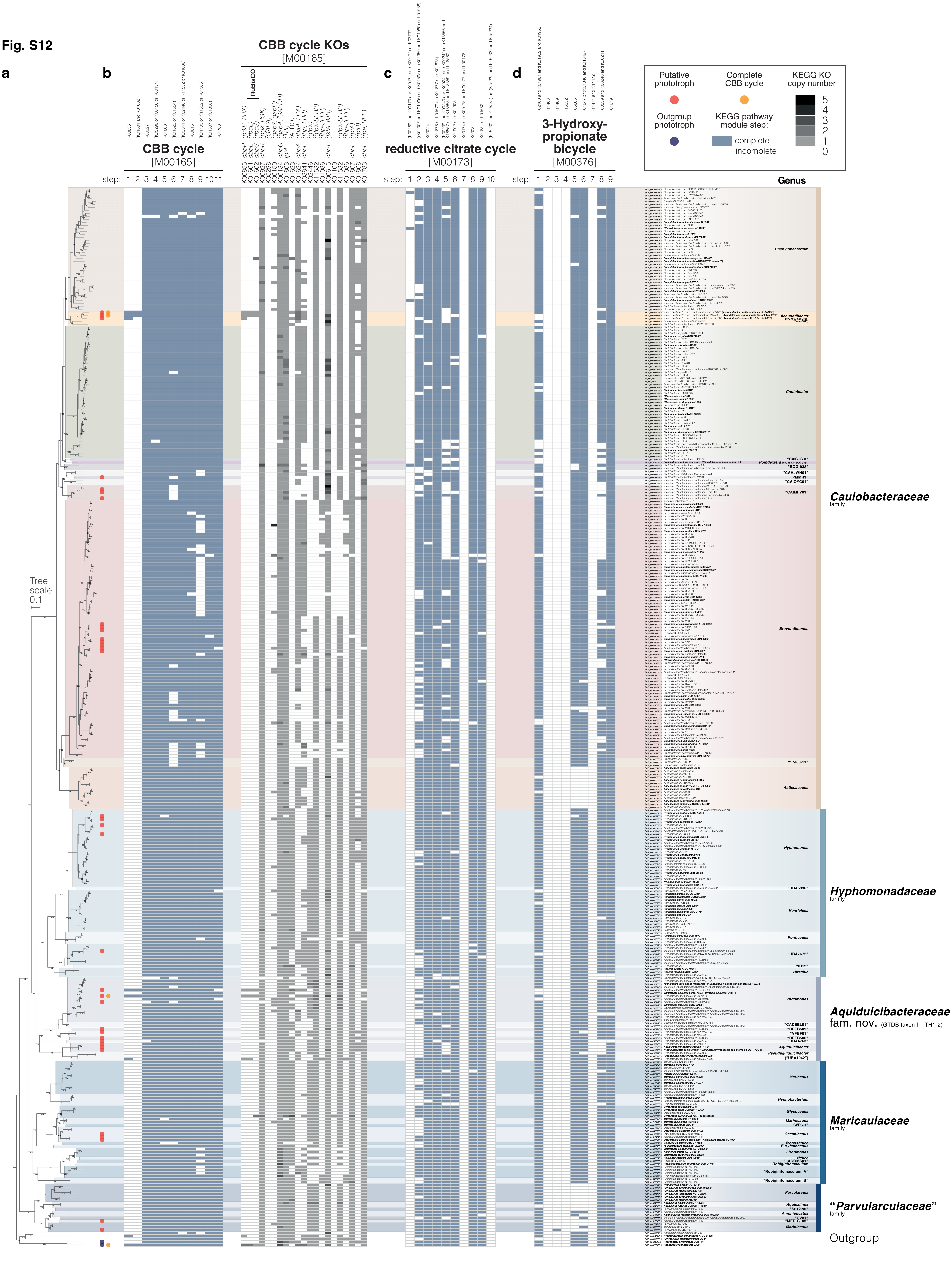
a
b
c
d
CBB cycle
[M00165]
CBB cycle KOs
[M00165]
RuBisCO
reductive citrate cycle
[M00173]
3-Hydroxy-propionate bicycle
[M00376]
step:
Putative phototroph
Complete CBB cycle
KEGG KO copy number
Outgroup phototroph
KEGG pathway module step:
complete
incomplete
Genus
Tree scale
0.1
Phenylobacterium
Acaudatibacter
Caulobacter
Poindextera
Brevundimonas
Asticcacaulis
Hyphomonas
Henriciella
Ponticaulis
Hirschia
Vitreimonas
Aquidulcibacter
Pseudaquidulcibacter
Maricaulis
Hyphobacterium
Glycocaulis
Marinicauda
Oceanicaulis
Woodsholea
Euryhalocaulis
Litorimonas
Hellea
Robiginitomaculum
Parvularcula
Aquisalinus
Amphiplicatus
Caulobacteraceae
family
Hyphomonadaceae
family
Aquidulcibacteraceae
fam. nov. (GTDB taxon f__TH1-2)
Maricaulaceae
family
"Parvularculaceae"
family
Outgroup

**Fig. S13**

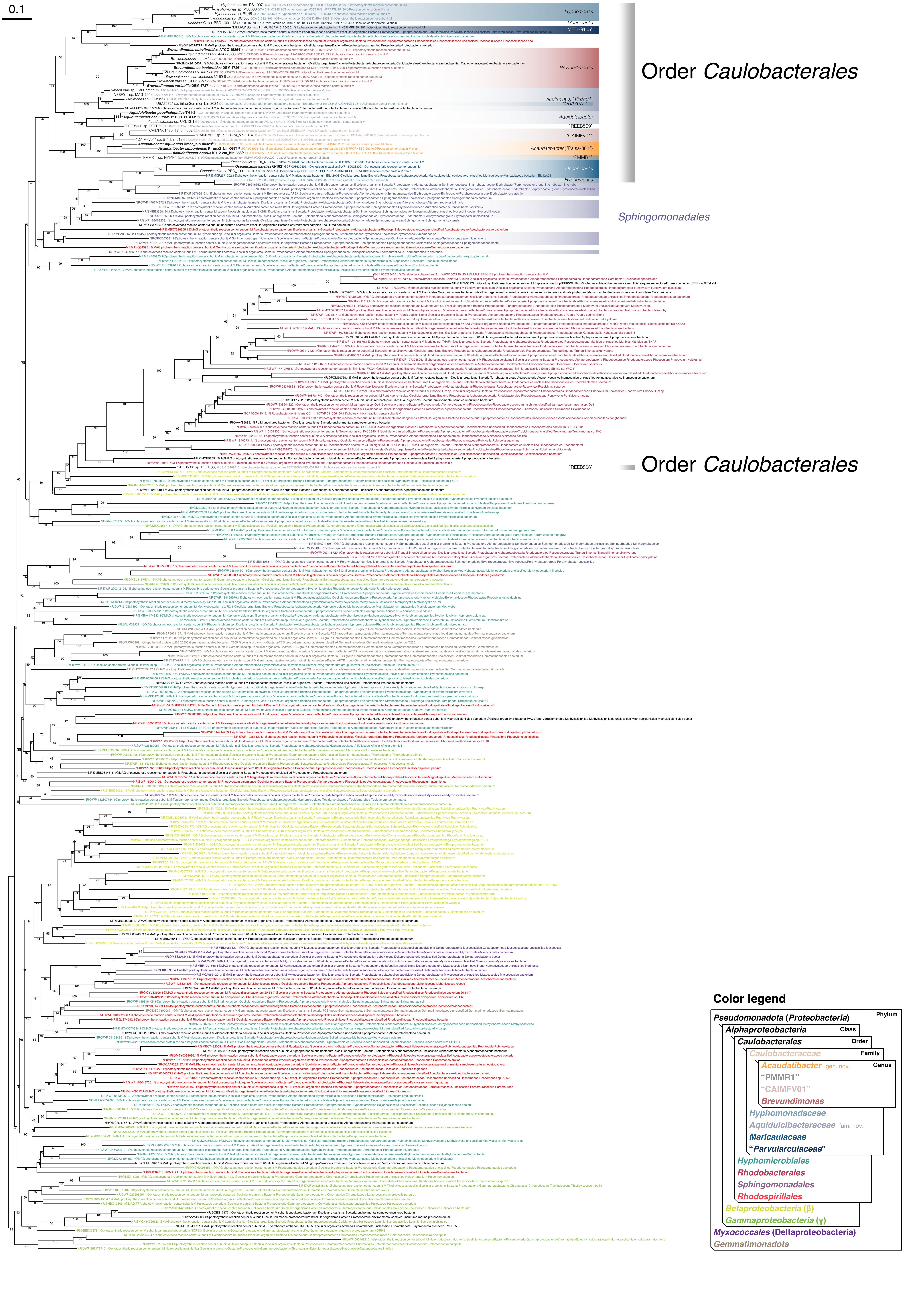
0.1
Hyphomonas
Marinicaulis
"MED-G105"
Brevundimonas
Vitreimonas, "VFBF01"
"UBA7672"
Aquidulcibacter
"REEB509"
"CAIMFV01"
Acaudatibacter ("Paisa-881")
"PMMR1"
Oceanicaulis
Hyphomonas
Order Caulobacterales
Sphingomonadales
"REEB506"
Order Caulobacterales
Color legend
Pseudomonadota (Proteobacteria)
Phylum
Alphaproteobacteria
Class
Caulobacterales
Order
Caulobacteraceae
Family
Acaudatibacter gen. nov.
Genus
"PMMR1"
"CAIMFV01"
Brevundimonas
Hyphomonadaceae
Aquidulcibacteraceae fam. nov.
Maricaulaceae
"Parvularculaceae"
Hyphomicrobiales
Rhodobacterales
Sphingomonadales
Rhodospirillales
Betaproteobacteria (β)
Gammaproteobacteria (γ)
Myxococcales (Deltaproteobacteria)
Gemmatimonadota

#### Fig. S14

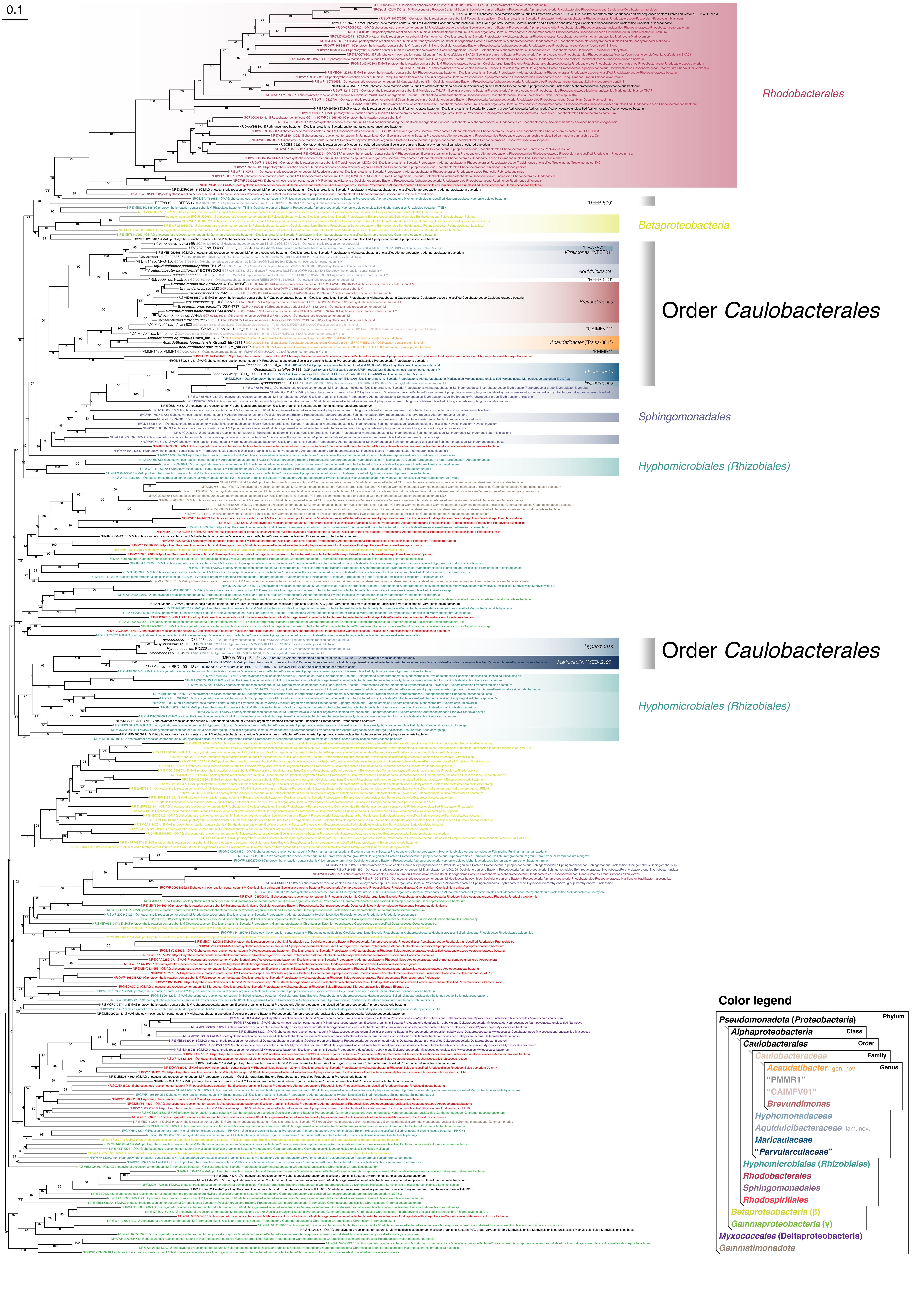
0.1
Rhodobacterales
Betaproteobacteria
Order Caulobacterales
Sphingomonadales
Hyphomicrobiales (Rhizobiales)
Order Caulobacterales
Hyphomicrobiales (Rhizobiales)
Color legend
Phylum
Pseudomonadota (Proteobacteria)
Class
Alphaproteobacteria
Order
Caulobacterales
Family
Caulobacteraceae
Genus
Acaudatibacter gen. nov.
"PMMR1"
"CAIMFV01"
Brevundimonas
Hyphomonadaceae
Aquidulcibacteraceae fam. nov.
Maricaulaceae
"Parvularculaceae"
Hyphomicrobiales (Rhizobiales)
Rhodobacterales
Sphingomonadales
Rhodospirillales
Betaproteobacteria (β)
Gammaproteobacteria (γ)
Myxococcales (Deltaproteobacteria)
Gemmatimonadota

### Fig. S15

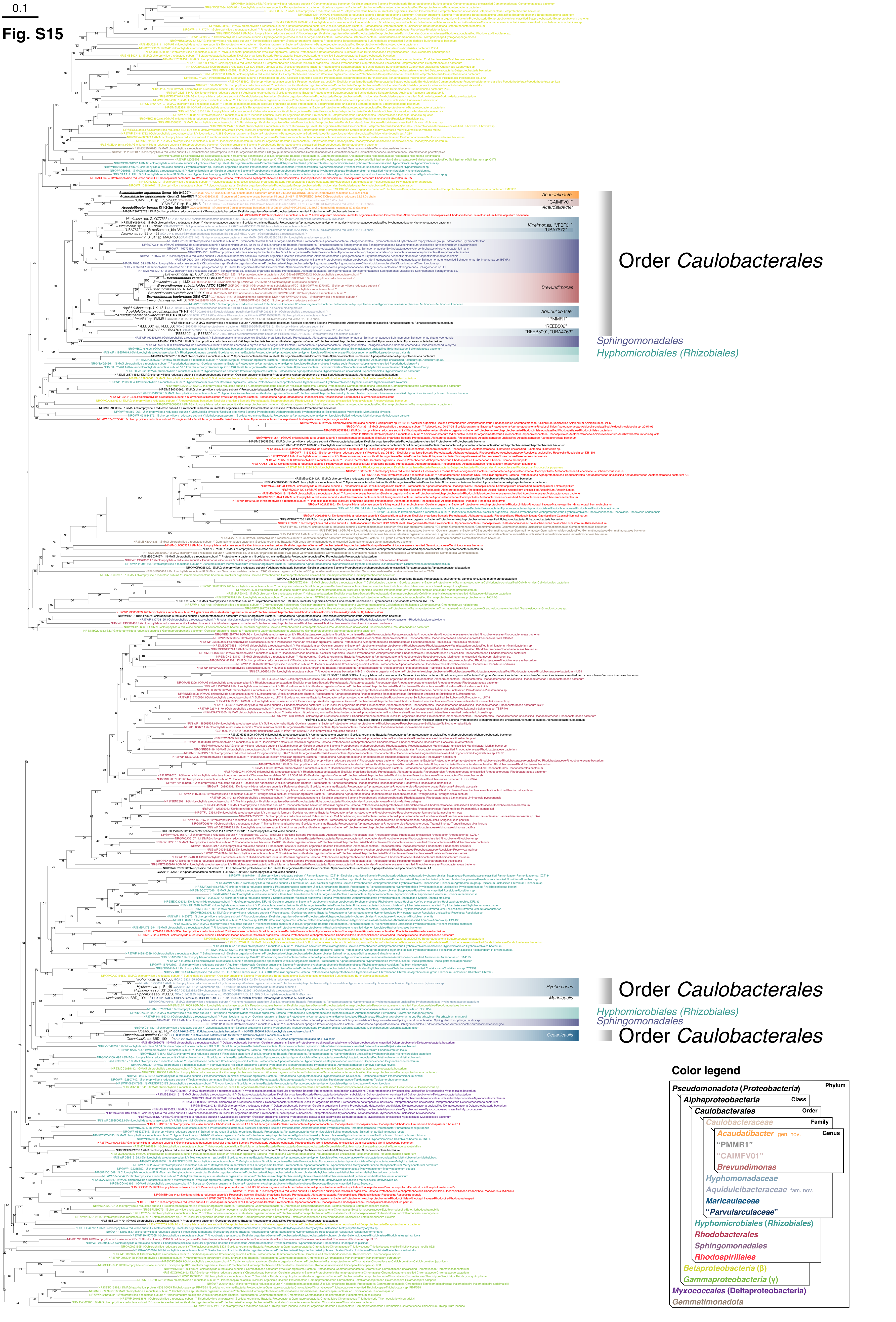
0.1
Acaudatibacter
"CAIMFV01"
Acaudatibacter
Vitreimonas, "VFBF01"
"UBA7672"
Order Caulobacterales
Brevundimonas
Aquidulcibacter
"PMMR1"
"REEB506"
"REEB509", "UBA4763"
Sphingomonadales
Hyphomicrobiales (Rhizobiales)
Hyphomonas
Order Caulobacterales
Maricaulis
Hyphomicrobiales (Rhizobiales)
Sphingomonadales
Order Caulobacterales
Oceanicaulis
Color legend
Pseudomonadota (Proteobacteria)
Phylum
Alphaproteobacteria
Class
Caulobacterales
Order
Caulobacteraceae
Family
Acaudatibacter gen. nov.
Genus
"PMMR1"
"CAIMFV01"
Brevundimonas
Hyphomonadaceae
Aquidulcibacteraceae fam. nov.
Maricaulaceae
"Parvularculaceae"
Hyphomicrobiales (Rhizobiales)
Rhodobacterales
Sphingomonadales
Rhodospirillales
Betaproteobacteria (β)
Gammaproteobacteria (γ)
Myxococcales (Deltaproteobacteria)
Gemmatimonadota

### Fig. S16

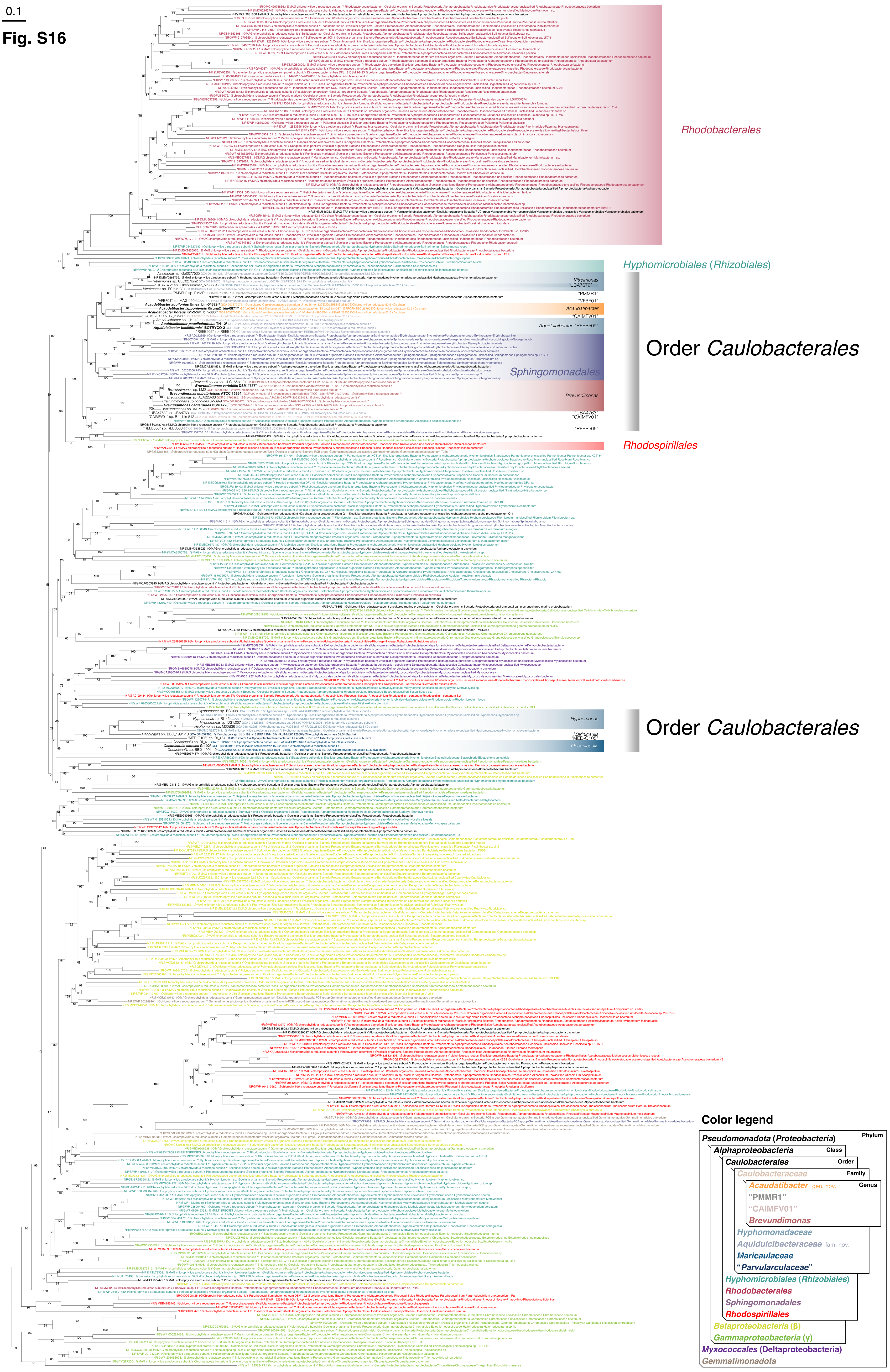
0.1
Rhodobacterales
Hyphomicrobiales (Rhizobiales)
Vitreimonas
"UBA7672"
"PMMR1"
"VFBF01"
Acaudatibacter
"CAIMFV01"
Aquidulcibacter, "REEB509"
Order Caulobacterales
Sphingomonadales
Brevundimonas
"UBA4763"
"CAIMFV01"
"REEB506"
Rhodospirillales
Hyphomonas
Maricaulis
"MED-G105"
Oceanicaulis
Order Caulobacterales
Color legend
Phylum
Pseudomonadota (Proteobacteria)
Class
Alphaproteobacteria
Order
Caulobacterales
Family
Caulobacteraceae
Genus
Acaudatibacter gen. nov.
"PMMR1"
"CAIMFV01"
Brevundimonas
Hyphomonadaceae
Aquidulcibacteraceae fam. nov.
Maricaulaceae
"Parvularculaceae"
Hyphomicrobiales (Rhizobiales)
Rhodobacterales
Sphingomonadales
Rhodospirillales
Betaproteobacteria (β)
Gammaproteobacteria (γ)
Myxococcales (Deltaproteobacteria)
Gemmatimonadota

### Fig. S17

**a**

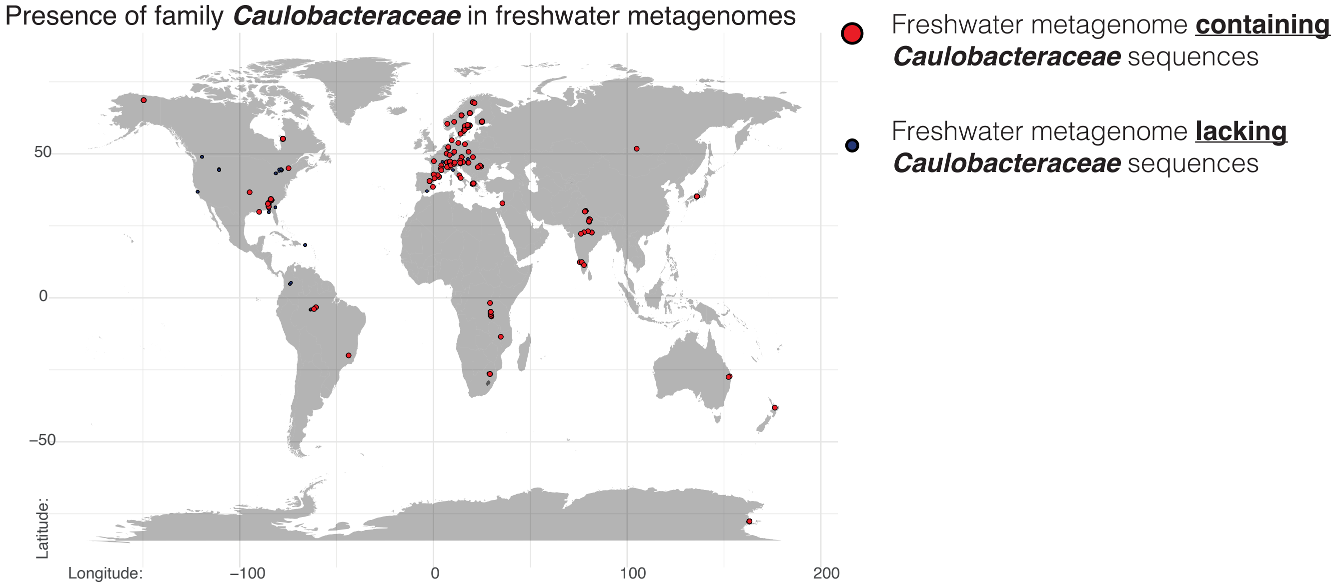
Presence of family Caulobacteraceae in freshwater metagenomes
Freshwater metagenome containing Caulobacteraceae sequences
Freshwater metagenome lacking Caulobacteraceae sequences
50
0
−50
Latitude:
Longitude:
−100
0
100
200

**b**

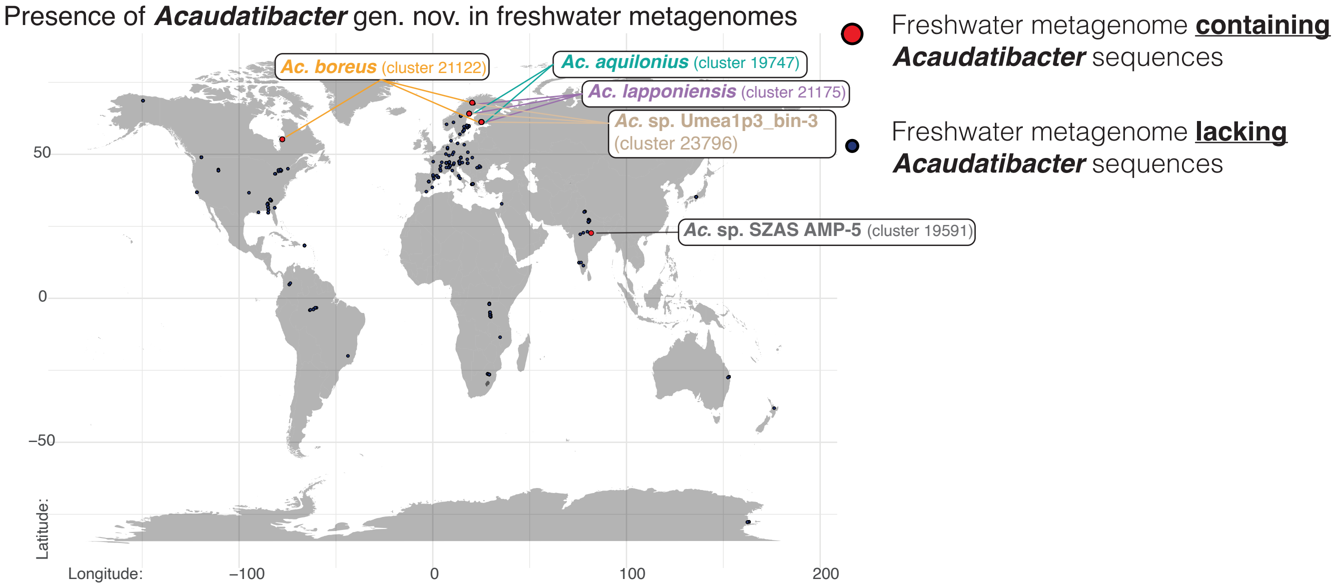
Presence of Acaudatibacter gen. nov. in freshwater metagenomes
Ac. boreus (cluster 21122)
Ac. aquilonius (cluster 19747)
Ac. lapponiensis (cluster 21175)
Ac. sp. Umea1p3_bin-3 (cluster 23796)
Ac. sp. SZAS AMP-5 (cluster 19591)
Freshwater metagenome containing Acaudatibacter sequences
Freshwater metagenome lacking Acaudatibacter sequences
50
0
−50
Latitude:
Longitude:
−100
0
100
200

**C**

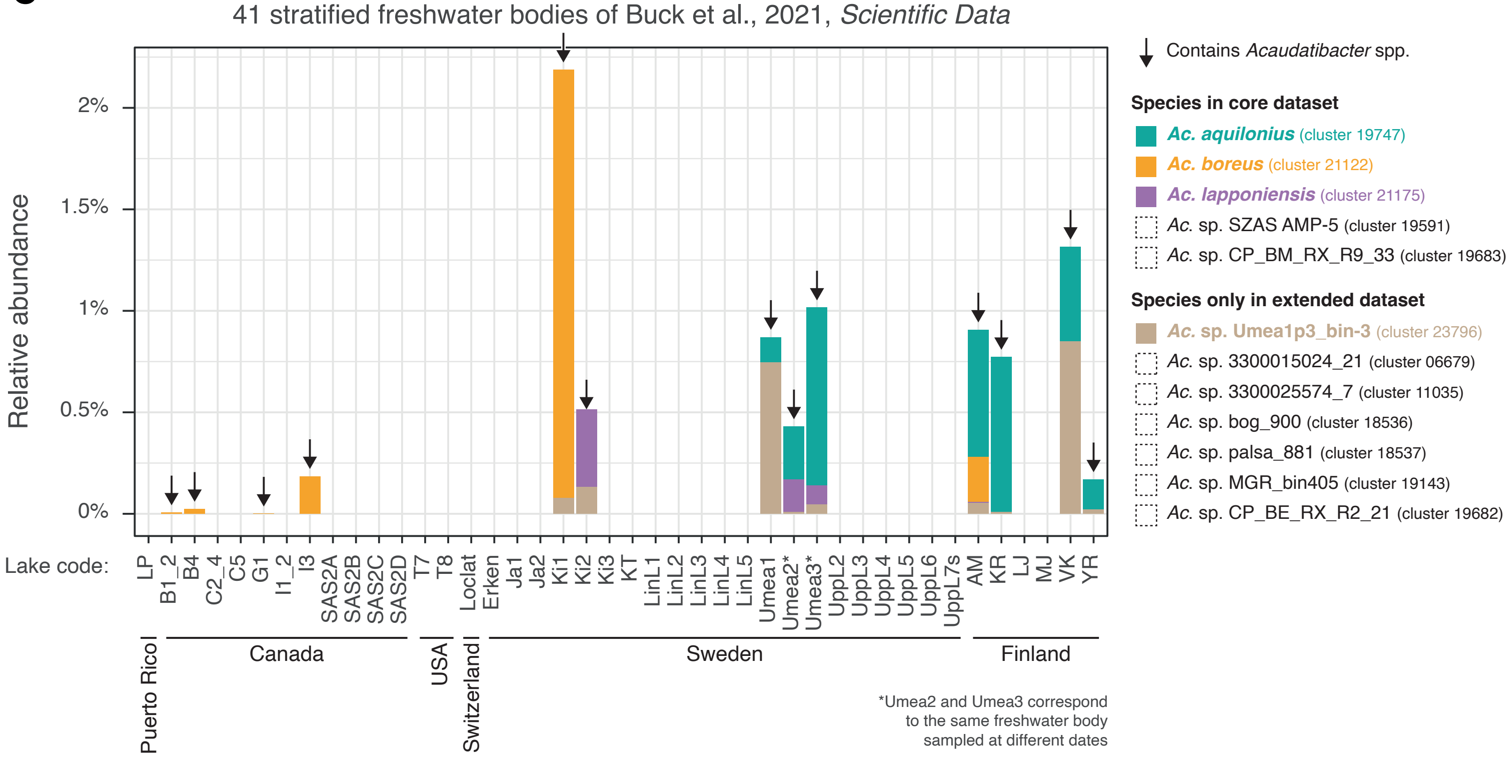
41 stratified freshwater bodies of Buck et al., 2021, Scientific Data
Relative abundance
2%
1.5%
1%
0.5%
0%
Contains Acaudatibacter spp.
Species in core dataset
Ac. aquilonius (cluster 19747)
Ac. boreus (cluster 21122)
Ac. lapponiensis (cluster 21175)
Ac. sp. SZAS AMP-5 (cluster 19591)
Ac. sp. CP_BM_RX_R9_33 (cluster 19683)
Species only in extended dataset
Ac. sp. Umea1p3_bin-3 (cluster 23796)
Ac. sp. 3300015024_21 (cluster 06679)
Ac. sp. 3300025574_7 (cluster 11035)
Ac. sp. bog_900 (cluster 18536)
Ac. sp. palsa_881 (cluster 18537)
Ac. sp. MGR_bin405 (cluster 19143)
Ac. sp. CP_BE_RX_R2_21 (cluster 19682)
Lake code:
LP
B1_2
B4
C2_4
C5
G1
I1_2
I3
SAS2A
SAS2B
SAS2C
SAS2D
T7
T8
Loclat
Erken
Ja1
Ja2
Ki1
Ki2
Ki3
KT
LinL1
LinL2
LinL3
LinL4
LinL5
Umea1
Umea2*
Umea3*
UppL2
UppL3
UppL4
UppL5
UppL6
UppL7s
AM
KR
LJ
MJ
VK
YR
Puerto Rico
Canada
USA
Switzerland
Sweden
Finland
*Umea2 and Umea3 correspond to the same freshwater body sampled at different dates

### Fig. S18

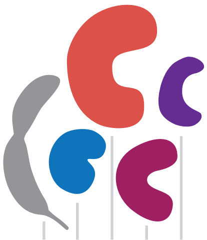

*Caulobacter crescentus* CB15 — GCF_000006905.1 — ***Caulobacterales*** order

*Chelatococcus reniformis* CGMCC 1.12919$^{T}$ — GCF_014640075.1 — ***Hyphomicrobiales*** order

*Roseiarcus fermentans* DSM 24875$^{T}$ — GCF_003315135.1

*Methylocystis* sp. Rockwell — GCF_000188155.2

*Methylocystis parva* OBBP$^{T}$ — GCF_027571405.1

| KEGG KO | Gene | *C. crescentus* CB15 | *C. reniformis* | *R. fermentans* | *Methylocystis* sp. Rockwell | *M. parva* | KEGG KO Annotation |
|---|---|---|---|---|---|---|---|
| K18642 | *creS* | ■ | ■ | 2 | ■ | ■ | crescentin |

**flagellar motility genes**

| KEGG KO | Gene | *C. crescentus* CB15 | *C. reniformis* | *R. fermentans* | *Methylocystis* sp. Rockwell | *M. parva* | KEGG KO Annotation |
|---|---|---|---|---|---|---|---|
| K02557 | *motB* | ■ | ■ | ■ | 2 | ■ | chemotaxis protein MotB |
| K13924 | *cheBR* | ■ | 2 | 2 | | ■ | two-component system, chemotaxis family, CheB/CheR fusion protein |
| K00575 | *cheR* | 3 | | 2 | 3 | 2 | chemotaxis protein methyltransferase CheR |
| K03407 | *cheA* | 4 | | | 4 | ■ | two-component system, chemotaxis family, sensor kinase CheA |
| K03412 | *cheB* | 2 | | | 2 | ■ | two-component system, chemotaxis family, protein-glutamate methylesterase/glutaminase |
| K03408 | *cheW* | 4 | | | 4 | ■ | purine-binding chemotaxis protein CheW |
| K03413 | *cheY* | 6 | | | 4 | ■ | two-component system, chemotaxis family, chemotaxis protein CheY |
| K02386 | *flgA* | ■ | | | ■ | ■ | flagellar basal body P-ring formation protein FlgA |
| K02387 | *flgB* | ■ | | | ■ | ■ | flagellar basal-body rod protein FlgB |
| K02388 | *flgC* | ■ | | | 2 | ■ | flagellar basal-body rod protein FlgC |
| K02389 | *flgD* | ■ | | | ■ | ■ | flagellar basal-body rod modification protein FlgD |
| K02390 | *flgE* | ■ | | | 2 | ■ | flagellar hook protein FlgE |
| K02391 | *flgF* | ■ | | | ■ | ■ | flagellar basal-body rod protein FlgF |
| K02392 | *flgG* | ■ | | | 2 | 2 | flagellar basal-body rod protein FlgG |
| K02393 | *flgH* | ■ | | | ■ | ■ | flagellar L-ring protein FlgH |
| K02394 | *flgI* | ■ | | | ■ | ■ | flagellar P-ring protein FlgI |
| K02396 | *flgK* | ■ | | | 2 | ■ | flagellar hook-associated protein 1 |
| K02397 | *flgL* | ■ | | | ■ | ■ | flagellar hook-associated protein 3 FlgL |
| K02400 | *flhA* | ■ | | | ■ | ■ | flagellar biosynthesis protein FlhA |
| K02401 | *flhB* | ■ | | | ■ | ■ | flagellar biosynthesis protein FlhB |
| K02408 | *fliE* | ■ | | | ■ | ■ | flagellar hook-basal body complex protein FliE |
| K02409 | *fliF* | ■ | | | ■ | ■ | flagellar M-ring protein FliF |
| K02410 | *fliG* | ■ | | | ■ | ■ | flagellar motor switch protein FliG |
| K02412 | *fliI* | ■ | | | ■ | ■ | flagellum-specific ATP synthase |
| K02415 | *fliL* | ■ | | | ■ | ■ | flagellar protein FliL |
| K02416 | *fliM* | ■ | | | ■ | ■ | flagellar motor switch protein FliM |
| K02417 | *fliN* | 2 | | | ■ | ■ | flagellar motor switch protein FliN |
| K02419 | *fliP* | ■ | | | ■ | ■ | flagellar biosynthesis protein FliP |
| K02420 | *fliQ* | ■ | | | ■ | ■ | flagellar biosynthesis protein FliQ |
| K02421 | *fliR* | ■ | | | ■ | ■ | flagellar biosynthesis protein FliR |
| K03406 | *mcp* | 16 | | | 8 | ■ | methyl-accepting chemotaxis protein |
| K02556 | *motA* | ■ | | | ■ | ■ | chemotaxis protein MotA |
| K03411 | *cheD* | ■ | | | ■ | | chemotaxis protein CheD |
| K03409 | *cheX* | ■ | | | ■ | | chemotaxis protein CheX |
| K10564 | *motC* | | | | ■ | ■ | chemotaxis protein MotC |
| K02411 | *fliH* | ■ | | | | | flagellar assembly protein FliH |
| K02413 | *fliJ* | ■ | | | | | flagellar protein FliJ |
| K02418 | *fliO, fliZ* | ■ | | | | | flagellar protein FliO/FliZ |
| K10565 | *motD* | | | | | ■ | chemotaxis protein MotD |
| K03410 | *cheC* | | | | | | chemotaxis protein CheC |
| K03415 | *cheV* | | | | | | two-component system, chemotaxis family, chemotaxis protein CheV |
| K03414 | *cheZ* | | | | | | chemotaxis protein CheZ |
| K02395 | *flgJ* | | | | | | peptidoglycan hydrolase FlgJ |
| K02399 | *flgN* | | | | | | flagellar biosynthesis protein FlgN |
| K24344 | *flgO* | | | | | | flagellar H-ring protein FlgO |
| K09860 | *flgP* | | | | | | outer membrane protein FlgP |
| K24346 | *flgQ* | | | | | | flagellar motility protein FlgQ |
| K24343 | *flgT* | | | | | | flagellar H-ring protein FlgT |
| K03516 | *flhE* | | | | | | flagellar protein FlhE |
| K02407 | *fliD* | | | | | | flagellar hook-associated protein 2 |
| K02414 | *fliK* | | | | | | flagellar hook-length control protein FliK |
| K13820 | *fliR-flhB* | | | | | | flagellar biosynthesis protein FliR/FlhB |
| K02422 | *fliS* | | | | | | flagellar secretion chaperone FliS |
| K02423 | *fliT* | | | | | | flagellar protein FliT |
| K21217 | *motX* | | | | | | sodium-type polar flagellar protein MotX |
| K21218 | *motY* | | | | | | sodium-type flagellar protein MotY |

tight-adherence type IV **pilus** genes

| KEGG KO | Gene | *C. crescentus* CB15 | *C. reniformis* | *R. fermentans* | *Methylocystis* sp. Rockwell | *M. parva* | KEGG KO Annotation |
|---|---|---|---|---|---|---|---|
| K02651 | *flp, pilA* | 2 | | ■ | 4 | 4 | pilus assembly protein Flp/PilA |
| K02278 | *cpaA, tadV* | ■ | | | ■ | ■ | prepilin peptidase CpaA |
| K02279 | *cpaB, rcpC* | ■ | | | ■ | ■ | pilus assembly protein CpaB |
| K02280 | *cpaC, rcpA* | ■ | | | ■ | ■ | pilus assembly protein CpaC |
| K02281 | *cpaD* | ■ | | | ■ | ■ | pilus assembly protein CpaD |
| K02282 | *cpaE, tadZ* | ■ | | | ■ | ■ | pilus assembly protein CpaE |
| K02283 | *cpaF, tadA* | ■ | | | 3 | ■ | pilus assembly protein CpaF |
| K12510 | *tadB* | ■ | | | ■ | ■ | tight adherence protein B |
| K12511 | *tadC* | ■ | | | ■ | ■ | tight adherence protein C |
| K12512 | *tadD* | | | | | | tight adherence protein D |
| K12513 | *tadE* | | | | | | tight adherence protein E |
| K12514 | *tadF* | | | | | | tight adherence protein F |
| K12515 | *tadG* | | | | | | tight adherence protein G |
